# SR2P: an efficient stacking method to predict protein abundance from gene expression in spatial transcriptomics data

**DOI:** 10.64898/2026.03.04.709692

**Authors:** Qingyue Wang, Anqi Gao, Yuying Li, Parth Khatri, Rong Hu, Jian Huang, Yudi Pawitan, Trung Nghia Vu, Huy Q. Dinh

## Abstract

Spatial transcriptomics data are largely available with RNA expression alone, limiting the detection of cell states defined by surface protein abundance. The lack of multi-omics spatial data limits the ability to identify immune cells and their signaling in the tumor microenvironment, as most solid tumors are immunologically poor and exhibit protein-RNA abundance discordance in critical immune cell surface markers. Although emerging technologies enable spatial multi-omics profiling, technical and cost constraints remain a hurdle. We introduce SR2P, a stacking-based machine-learning framework for predicting spatial protein abundance from RNA expression. SR2P integrates 11 complementary predictive models and consistently outperforms existing methods across multiple spatial multi-omics benchmark. We showcased an application of SR2P recovered macrophage-enriched regions and identified potential immune markers associated with therapeutic response from head-and-neck squamous cell carcinoma patients. SR2P enables protein-abundance inference from RNA-only spatial data, extending the analytical capabilities of current spatial platforms for studies of tumor immunology.

## Introduction

Proteins are the functional executors in cells that more directly reflect biological activity than RNA, and often the primary therapeutic targets ^1, 2^. Their spatial distribution determines cellular roles and interactions within the tissue microenvironment. At the same time, abnormal protein expression or localization serves as a key biomarker for disease subtype classification, immune profiling, and therapeutic stratification ^3–6^. Recent advances in spatial transcriptomics technologies, such as 10x Genomics Visium, Slide-seq, and Stereo-seq, have enabled genome-wide RNA profiling while preserving spatial organization within tissue sections. These methods have substantially advanced our understanding of tissue architecture, cell–cell communication, and spatial gene expression programs. However, spatial RNA-seq measurements alone provide only an indirect view of cellular function, as RNA abundance does not fully capture post-transcriptional regulation or protein-level activity. Spatial proteomics technologies, such as mass spectrometry-based methods ^7^ and CODEX ^8^, enable direct in situ mapping of protein abundance, but these methods remain costly, technically demanding, and low-throughput. As a result, most current spatial omics datasets contain only transcriptomic measurements, leaving protein-level insights inaccessible. Recent advances in spatial technologies enable us to profile both transcriptomics and proteomics simultaneously ^9–12^, including DBiT-seq and Spatial CITE-seq ^13, 14^, as well as antibody-assisted extensions of spatial transcriptomics platforms such as 10x Genomics Visium CytAssist ^14^, where targeted protein markers are measured in addition to spatial RNA expression. While they are not yet widely adopted, they offer an opportunity to evaluate and test the transferability of RNA and protein measurements across the same spatial regions.

Predicting protein abundance from RNA expression provides a practical, cost-effective strategy to expand the utility of spatial datasets. Such an approach can enable retrospective protein profiling in existing RNA-only spatial datasets and facilitate integrative multi-omics analyses. In particular, surface protein markers are a cornerstone of immunology research and essential for sorting specific immune subsets for further functional studies. However, the RNA–protein relationship is complex and influenced by multiple regulatory layers, including transcriptional and translational regulation, post-translational modifications, and protein degradation, leading to substantial RNA–protein mismatches^15,16^. Spatial heterogeneity further complicates this relationship, as protein abundance is modulated by tissue microenvironment and cell–cell interactions^17–19^. Finally, sequencing-based technologies such as Visium are often constrained by sparse spot-level RNA expression and protein abundance measurements, which are susceptible to stochastic noise and high variability, posing a major challenge for predictive modeling. A recent study proposes sclinear, a linear regression–based model for predicting protein abundance in single-cell RNA-seq datasets ^20,21^. While this method is theoretically applicable to spatial datasets, its performance in a spatial context has not yet been evaluated. Graph neural networks (GNNs) have emerged as powerful tools for spatial omics analysis because they naturally encode spatial relationships through graph-based representations^22–24^. By modeling spatial spots as nodes connected by proximity-based edges, architectures such as Graph Attention Networks (GAT), GraphSAGE can integrate transcriptomic profiles with spatial context. In particular, Dual Graph Attention Networks (DGAT)^25^ extend the conventional attention mechanism by jointly leveraging a feature graph and a spatial graph, enabling prediction of protein abundance from RNA expression in spatial CITE-Seq data.

Here, we developed SR2P, a stacking-based machine learning framework for spatial protein abundance prediction from transcriptomic data, and provided a systematic evaluation of its performance across multiple Visium CytAssist spatial CITE-seq datasets. SR2P integrated predictions from diverse base learners to utilize their complementary strengths under different spatial and biological contexts and was able to recover immune-enriched regions within tumors from head and neck squamous cell carcinoma (HNSCC) patients, thereby identifying potential protein markers associated with anti-PD-1 immune checkpoint blockade therapy response.

## Results

### SR2P - a stacking machine learning framework for spatial protein prediction

We hypothesized that different predictive models exploit distinct structural and gene expression resolutions in the complex and high-dimensional data, such as spatial transcriptomics data. Therefore, we developed SR2P framework (**Fig. 1a**, and **Suppl Fig. 1**) for predicting spatial protein abundance from spatial transcriptomic profiles, taking advantages of eleven models, including graph-based representation with GNNs as well as scalability and predictive power in structured data with tree-based ensemble models, particularly gradient-boosting frameworks such as XGBoost, LightGBM, and CatBoost. The details of stacking framework of SR2P (**Suppl Fig. 1**) are referred to section Material and Methods. To comprehensively assess model performance, we benchmark SR2P against the eleven competing model configurations, including partial least squares regression (PLS) ^26^, three gradient-boosting ensemble models (XGBoost ^27^, LightGBM ^28^, and CatBoost ^29, 30^), and three GNN architectures (GAT ^31^, GraphSAGE ^32^, and DGAT ^25^), each evaluated with and without spatial neighborhood augmentation. Model performance is assessed under three biologically relevant evaluation settings: (1) within-sample validation, (2) within-tissue-type validation, and (3) cross-tissue validation with four distinct tissue types (**Fig. 1b**). This design enables us to quantify the impact of stacking-based integration and spatial modeling on protein abundance prediction and to evaluate the generality of predictive models trained on data from different tissue types.

**Figure 1:**
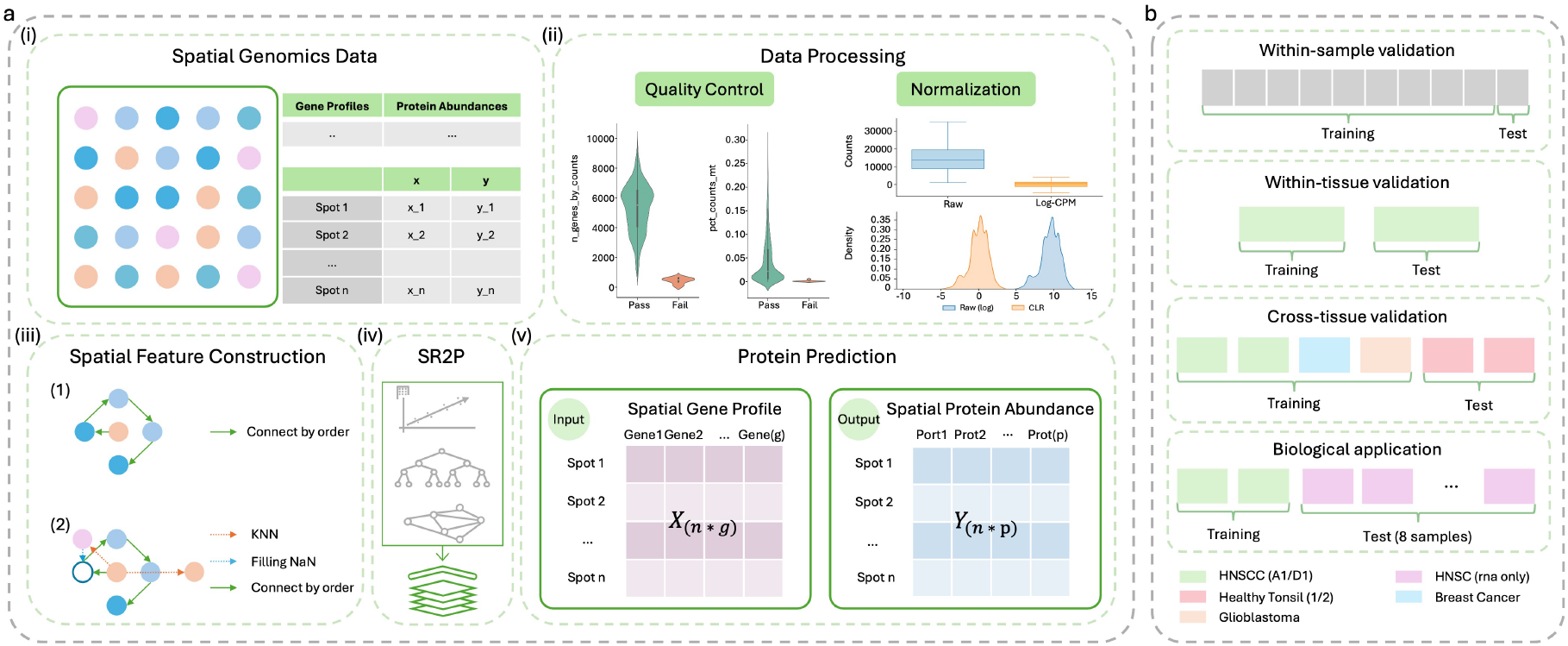
Overview of SR2P for spatial protein abundance prediction from spatial transcriptomics data. **a** Workflow for model construction and spatial protein prediction. **(i)** Spatial genomics data consist of paired spatial transcriptomic (gene expression) and spatial proteomic (protein abundance) measurements aligned at matched spatial coordinates (spots). **(ii)** Data preprocessing includes quality control to filter low-quality spots and normalization to reduce technical variability and scale differences. **(iii)** Spatial feature construction incorporates local tissue context using ordered neighborhood structures, k-nearest neighbor (KNN) relationships, and imputation for missing neighbors. **(iv)** SR2P utilizes the stacking approach, integrating the multiple predictive models as its base learners. The base learners include linear models, tree-based ensemble models, and graph-based models. **(v)** Each model takes a spatial gene expression matrix *X*_*n*×*g*_, where *n* is the number of spots and *g* is the number of genes, and predicts a spatial protein abundance matrix *Y*_*n*×*p*_, where *p* is the number of proteins. **b** Model performance is assessed using within-sample validation, within-tissue validation, and cross-tissue validation to evaluate predictive accuracy and generalization. Additionally, trained SR2P models are applied to biological case studies, in which spatial protein abundance is predicted from spatial transcriptomic data in independent samples. Colors indicate different tissue types, and grey indicates unspecified or any.

### Spatial information improves protein abundance prediction from RNA expression with within-sample cross-validation

To differentiate the same problem with scRNA-Seq, we evaluate the impact of spatial information in different predictive models using spatial 10-fold cross-validation with the within-sample validation setting (details in Section **Materials and Methods**). Briefly, k-means clustering is performed on the spatial coordinates of spots to construct 10 spatially coherent folds (**Suppl Fig. S2** shows the spatial layout of folds for each dataset). Overall, all methods were able to predict spatial protein abundance accurately, especially for the canonical markers for major cell type identification, such as CD45 (PTPRC) for immune cells, CD163 for macrophages, VIM for stromal fibroblast cells, and EPCAM for tumor/epithelia cells (representative example with the Breast Cancer dataset and four methods including SR2P, PLS-Spatial, CatBoost-Spatial, and DGAT, **Fig. 2a**). SR2P consistently produces spatially coherent predictions that closely match the observed protein distributions. CatBoost-Spatial also closely recapitulates the spatial patterns in the ground truth from observed data, especially for the CD163 marker, whereas DGAT yields more variable and spatially diffuse predictions. Incorporating spatial neighborhood features significantly improves performance for non-GNN models, especially in the HNSCC and tonsil samples (*p* < 0.05), underscoring the importance of spatial information (**Fig. 2b**). Among GNN models, DGAT achieves slightly higher median performance and lower variance than GAT and GraphSAGE, but still falls short of the spatially enhanced tree-based models, particularly LightGBM-Spatial, XGBoost-Spatial, and CatBoost-Spatial, which are consistently among the high-performing methods. Compared with spatially enhanced learning-based models, SR2P achieves superior performance across multiple datasets while maintaining low variance (**Suppl Figs. S3 and S4**).

**Figure 2:**
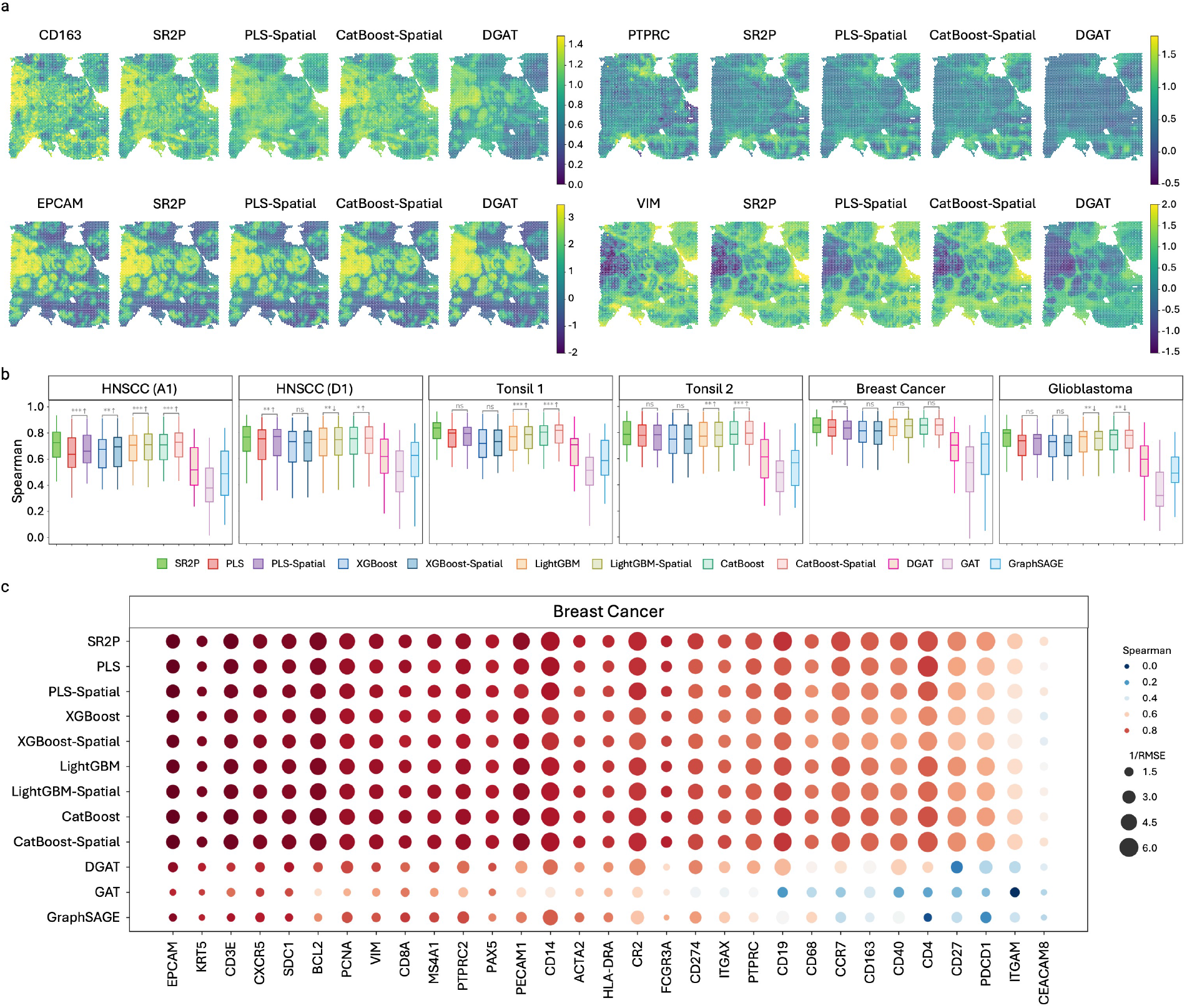
Comparison of protein prediction models across six spatial genomics datasets using spatial within-sample validation. **(a)** Spatial protein abundance maps for representative immune and tumor markers in the breast cancer dataset. For each protein, the measured abundance is shown alongside predictions from the top-performing models in each model class, enabling visual comparison of spatial localization patterns. Predicted maps are scaled to match the dynamic range of measured protein abundance for visual comparison. **(b)** Prediction accuracy of eleven models for each dataset, evaluated using Spearman correlation between predicted and measured protein abundances. The two-tailed Wilcoxon rank-sum test is used to assess differences between spatial and non-spatial settings for non-GNN methods (*p* < 0.05 (*), *p* < 0.01 (**), and *p* < 0.001 (***), with *ns* indicating no significant difference). Arrows indicate the direction of performance change, where upward (downward) arrows denote that the spatial setting performs better (worse) than the non-spatial setting. **(c)** Protein-wise prediction performance on the Breast Cancer dataset. Color indicates Spearman correlation, and dot size represents the inverse root mean squared error (1/RMSE), with larger dots indicating better performance.

Next, we evaluate the performance of predictive models for individual protein markers (**Fig. 2c**), SR2P achieves high accuracy for a subset of surface protein markers with strong expression and coherent spatial organization in breast cancer tissues, including epithelial markers (e.g., EPCAM, KRT5, SDC1), stromal cell markers (e.g., PECAM1, VIM), and immune-associated markers (e.g., CD3E, CD8A, PAX5), as reflected by higher Spearman correlations and lower RMSE values. In contrast, proteins with low abundance or weak spatial structure (e.g., myeloid cell markers like ITGAM/CD11b and CEACAM8/CD66b) are more challenging to predict across all models (Fig. 2c). Neutrophils (marked by CD66b protein expression) were known to be poorly captured in tumors, similar to our previous analysis with HNSCC^33^. Notably, the specific proteins best predicted vary across datasets and reflect the prevailing specificity of cell populations and tumor microenvironment structure in each sample. Similar tissue-specific predictive patterns are observed in the other tissue types (see **Suppl Fig. S5 & S6**).

### Tissue specificity in spatial protein abundance predictions from RNA gene expression

Molecular and cellular information, including protein and RNA gene expression, is tissue-specific ^34^ and controlled by a complex gene regulatory program. Taking advantage of the ML prediction frameworks, we evaluate the ability of the prediction models to generalize within and across tissue types. First, for the case where both training and testing are performed within the same tissue, we used (i) HNSCC tumor samples and their closely related normal tissue and (ii) Tonsil datasets, where most models achieve moderate to high protein-wise prediction performance (**Fig. 3a**). SR2P is consistently among the top performance against other methods for both HNSCC and Tonsil samples. In addition, spatially augmented tree-based ensemble models, including LightGBM-Spatial and CatBoost-Spatial, demonstrate strong, competitive performance. This indicates that spatial message passing alone in GNN models is insufficient to capture predictive signals without strong feature-level modeling. These findings reinforce the advantage of flexible boosting frameworks when combined with spatial context.

**Figure 3:**
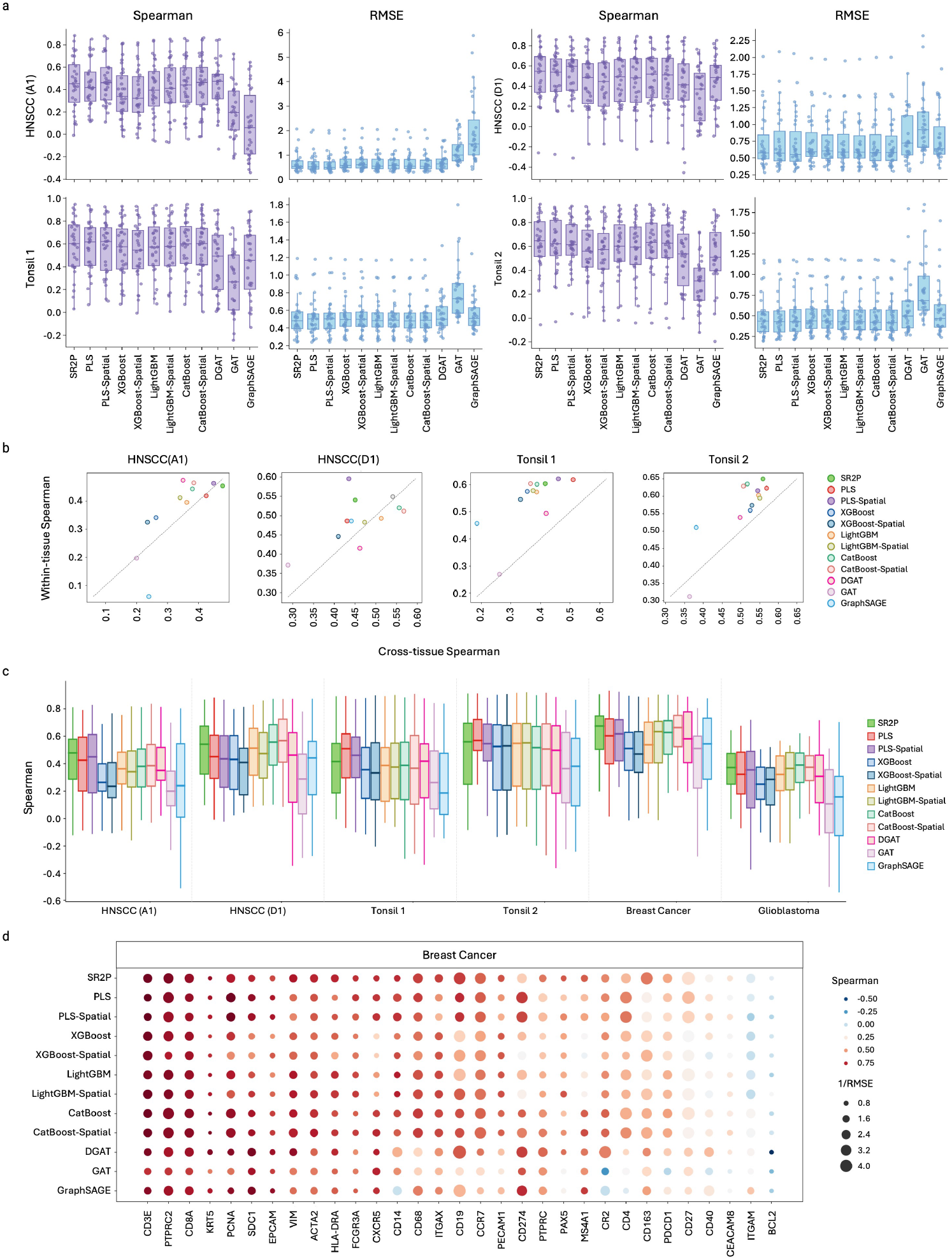
Model generalization within and across tissue types for spatial protein abundance prediction. **(a)** Within-tissue prediction performance for HNSCC and Tonsil datasets, shown as protein-wise Spearman correlation and RMSE for each model. **(b)** Cross-tissue prediction performance, where each tissue is evaluated as a held-out test dataset. Boxplots summarize the distribution of protein-wise Spearman correlations across models. **(c)** Protein-level cross-tissue prediction performance for the Breast Cancer dataset. Color denotes Spearman correlation, and dot size represents inverse RMSE (1/RMSE), with larger dots indicating better accuracy. **(d)** Comparison of within-tissue and cross-tissue performance across models. Each point represents the median Spearman correlation under the two validation settings, with deviations from the diagonal indicating loss of generalization across tissues.

In most models, we observed a consistent decrease in predictive accuracy when tissue type differed between training and testing data, compared with within-tissue training models (**Fig. 3b**). CatBoost-Spatial and LightGBM-Spatial exhibit the smallest performance gaps, while SR2P maintains the highest overall accuracy with only moderate performance degradation across tissues, whereas GNN-based models show the most significant drops, suggesting limited robustness to cross-tissue prediction. Next, we asked how well predictions will perform if models were trained on samples from different tissue types and applied to a held-out tissue type. Cross-tissue evaluation shows an apparent reduction in prediction accuracy compared to the within-tissue setting as median Spearman correlations decrease across nearly all methods (**Fig. 3c**), indicating limited transferability of protein–RNA spatial relationships across tissues.

SR2P remains highly competitive under this setting, consistently ranking among the top-performing methods across datasets, with median Spearman correlations of approximately *ρ* = 0.40–0.55 in HNSCC samples and *ρ* = 0.45–0.60 in breast cancer. In the tonsil datasets, spatially augmented tree-based ensemble models, particularly CatBoost-Spatial and LightGBM-Spatial, achieve the highest median performance, reaching correlations of approximately *ρ* = 0.50–0.60, whereas SR2P attains slightly lower but still comparable accuracy (typically *ρ* = 0.45–0.55). In contrast, GNN-based models, such as DGAT and GraphSAGE, show substantially lower cross-tissue performance, often below *ρ* = 0.30, with increased variability. As expected, performance varies notably across tissues: breast cancer and tonsil samples retain moderate predictive accuracy. These results indicate that spatial protein abundance models trained on one tissue type do not generalize uniformly across tissues, and prediction accuracy is strongly influenced by biological similarity between training and testing tissue contexts.

To evaluate how cross-tissue predictions perform at individual protein markers, similar to the previous anal-ysis (**Fig. 2c**), we found that predictive performance varies substantially across protein targets in the Breast Cancer dataset (**Figure 3d**). However, immune-associated markers such as CD3E, CD8 (T cells), PTPRC2 (CD45+ immune cells), and SDC1 (plasma/B cells) demonstrate relatively high correlations and low RMSE values across multiple models, indicating that their expression patterns are more conserved across tissues, as expected. In contrast, markers with stronger tissue-specific or tumor-specific expression profiles, such as EPCAM, show marked reductions in predictive accuracy, suggesting that the underlying RNA-protein relationships for these proteins do not transfer well across tissue contexts. Additionally, DGAT and GAT exhibit consistently weaker protein-level performance, with larger error variance across targets, whereas SR2P, CatBoost-Spatial and LightGBM-Spatial maintain more stable accuracy profiles. These results highlight that the reliability of cross-tissue predictions is influenced not only by model architecture but also by the extent to which protein expression patterns and their transcriptomic determinants are preserved across tissue types. Cross-tissue evaluation of the remaining datasets (**Suppl Fig. S7 & S8**) further supports these results. **Suppl Figs S9-S14** illustrate for each sample the spatial distributions of two representative proteins alongside their model predictions, enabling visualization of tissue-specific prediction accuracy.

In summary, we show that prediction performance is strongly tissue-dependent. Models trained within the same tissue achieve higher and more stable accuracy, while cross-tissue generalization is limited and varies across protein targets and tissue microenvironments. These findings suggest that, whenever possible, spatial protein prediction should be performed using training data derived from the same or closely related tissue contexts.

### Spatial protein predictions recover immune-rich regions in the HNSCC tumor microenvironment and identify protein markers associated with immunotherapy response

To demonstrate the applications of spatial protein prediction models, we used training data from two HNSCC Visium Spatial-CITESeq samples to predict protein abundance for 31 markers across 8 Visium spatial transcriptome (RNA alone) samples from a HNSCC patient cohort treated with ICB immunotherapy.

First, we asked whether predicted proteins help us identify previously missing regions due to the lack of surface protein markers. In a poorly immune-infiltrated sample (**Suppl Fig. 15** - with cell-type deconvolution maps), we used spatial clustering using three feature representations: RNA-only, predicted protein-only, and a combined RNA+protein feature space. The combined RNA+protein clustering identifies 9.7% more of macrophage-enriched spatial spots compared to the single data modality (**Fig. 4a-b**). Macrophage-enriched spots were de-fined as spatial locations with the highest macrophage proportion, as estimated by DECLUST deconvolution ^35^. Specifically, the combination clustering recovered clusters 16 and 29, which are identified as macrophage-rich regions only in RNA- and protein-only analyses, respectively, among the highly enriched macrophage regions (clusters 1, 10, 16, 27, and 29) based on deconvolution. These clusters show high correlation (Pearson r = 0.77) in macrophage proportions estimated using deconvolution and macrophage gene signatures (see **Materials and Methods** section) from scRNA-Seq references (**Fig. 4b**). Notably, we found that macrophage-enriched clusters identified in the RNA-only (Cluster 2) and protein-only (Cluster 5) settings are re-identified as clusters (29 & 10) with higher enrichment in macrophage protein marker, CD163, compared to the rest of the tissue when using the combined RNA+protein clustering (*p* = 1.2 × 10^−6^ vs. *p* = 8.2 × 10^−21^, and *p* = 3.3 × 10^−11^ vs. *p* = 9.6 × 10^−23^, **Fig. 4c**). These data confirmed the necessity of integrating complementary transcriptomic and proteomic information, preserving distinctions captured by each modality alone, and producing more spatially coherent macrophage-rich region boundaries, thereby making inferred tissue domains more biologically interpretable.

**Figure 4:**
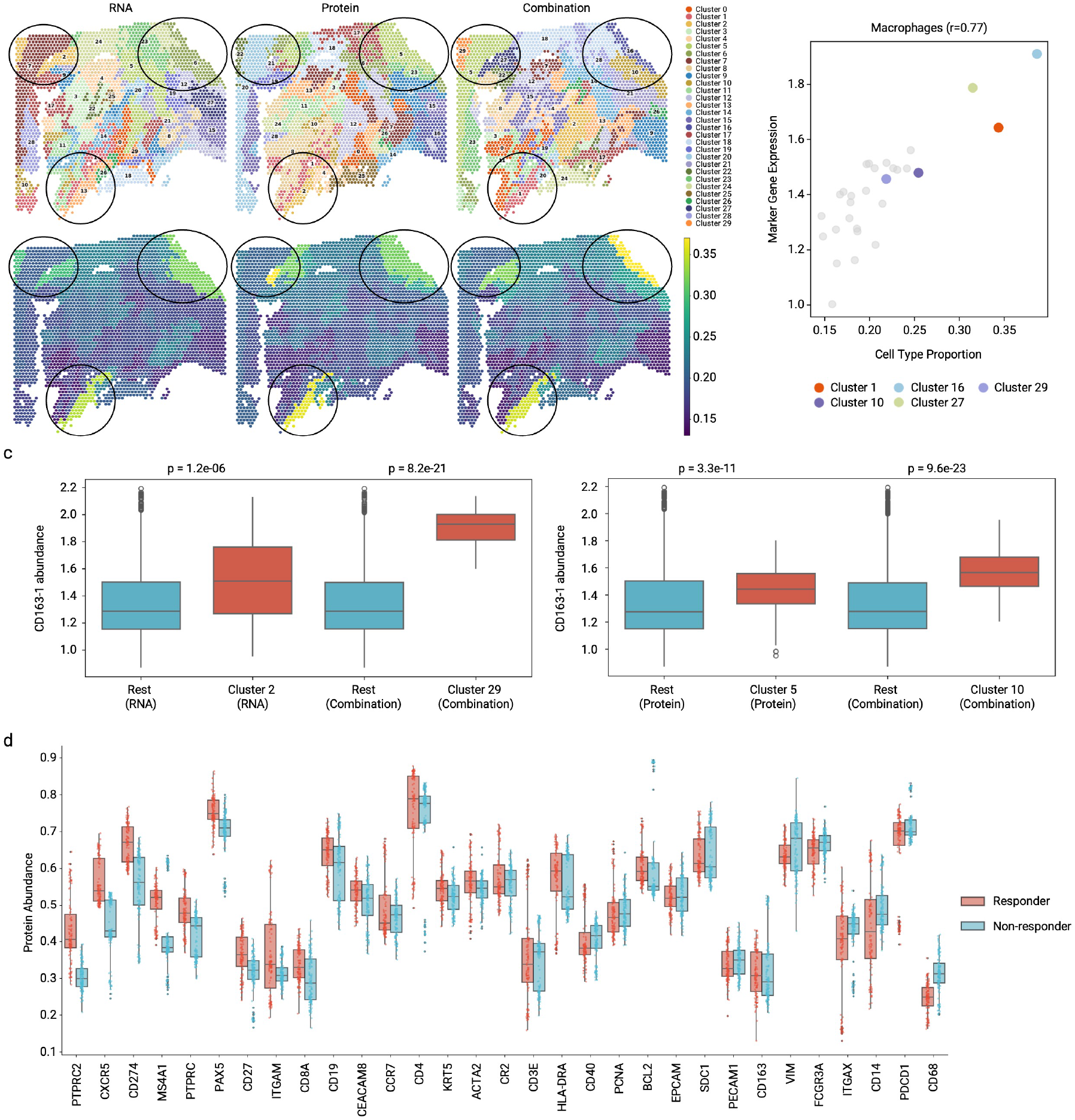
Spatial characterization of tumor immune microenvironment features using predicted protein abundance. **(a)** Spatial clustering of the HNSCC sample using RNA-only, protein-only, and combined RNA+protein features. The circled regions highlight macrophage-enriched spatial domains. **(b)** For the combined-feature clustering, the relationship between cluster-level macrophage cell-type proportion and the mean expression of macrophage-associated marker genes is shown, with each point representing a spatial cluster. **(c)** Differential CD163 abundance between macrophage-enriched clusters and the remaining tissue across RNA-based, protein-based, and combined feature clustering. **(d)** Comparison of predicted protein abundance between responder and non-responder groups across immune-related markers.

Next, we evaluated predicted protein markers that differ between Responder and Non-Responder patients using a combination of multi-omics and RNA-only Visium samples, resulting in a total of 10 samples. Given the limited number of patients, statistical comparisons were performed at the cluster level to increase analytical power (**Fig. 4d**; **Suppl Table S1**), yielding 116 clusters from Responder samples and 138 clusters from Non-Responder samples. We found that Responder-associated clusters exhibit higher known “immunologically hot tumor” phenotypes, including higher CD45+ markers (PTPRC and PTPRC2; adjusted *p* = 8.20 × 10^−14^ and 1.48 × 10^−30^), CD8+ T cell markers (CD8A; adjusted *p* = 1.29 × 10_−7_). In contrast, clusters in non-responder samples display elevated levels of macrophage- and myeloid-associated proteins, such as CD68, CD14, and IT-GAX, marking immunosuppressive myeloid infiltration phenotypes as observed in anti-PD1 resistance. These observed differences indicate the feasibility of using predicted protein information derived from RNA gene expression to identify proteomic immunotherapeutic biomarkers and targets.

### Computational time for protein abundance prediction

The computational efficiency of different models is compared under a cross-tissue validation setting, using the breast cancer dataset as the test set. All experiments are conducted on a Linux server equipped with 96 logical CPUs (48 physical cores) and no GPU acceleration. The reported computational cost corresponds to the test phase only, measured from the start of model inference in the cross-tissue validation until the results are exported. For the stacking based SR2P framework, the reported testing time reflects only the inference time of the meta learner, which is 2.19 seconds (**Fig. 5**), ranking as the third fastest among all methods. Of note, due to its stacking architecture, the overall inference time of SR2P also includes the cumulative inference time of its base learners. Among the remaining approaches, the graph based models GAT and GraphSAGE exhibit the lowest inference time of 0.37 and 0.57 seconds, respectively, demonstrating excellent computational efficiency. Tree-based models without spatial features, including XGBoost, LightGBM, and CatBoost, exhibit moderate computational time, ranging from 3.3 to 6.3 seconds. The incorporation of spatial features consistently increases the computational time across these models. In particular, LightGBM Spatial and CatBoost Spatial require 7.9 and 11.8 seconds, respectively. A similar pattern is observed for PLS, whose computational time increases substan-tially from 7.2 seconds to 26.1 seconds after spatial features are introduced. Overall, the results demonstrate that most models retain practical inference efficiency under the cross-tissue validation setting.

**Figure 5:**
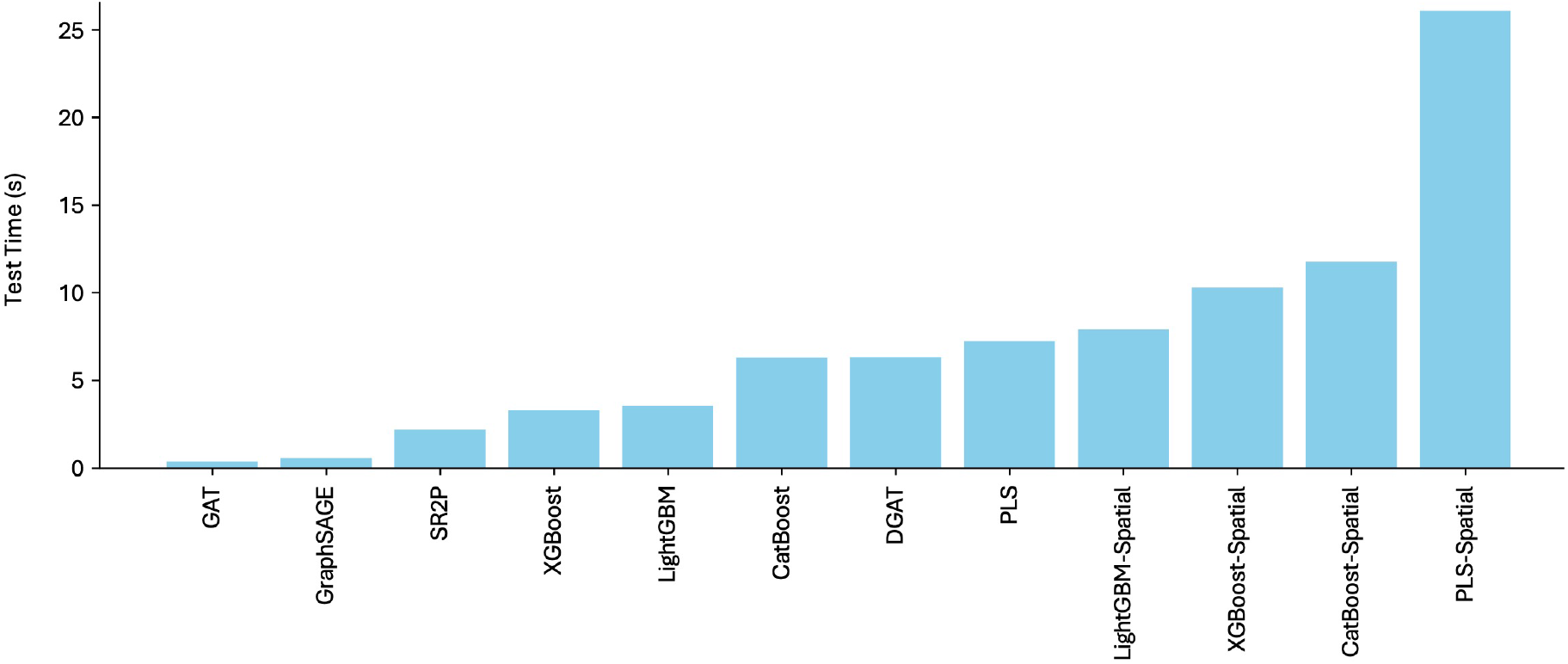
Computational time across 12 models in the cross-tissue validation. The x-axis represents the prediction models, while the y-axis shows the inference time in seconds.

## Discussion

This study introduces SR2P, an efficient stacking-based framework for predicting spatial protein abundance from spatial transcriptomic data. By integrating complementary ML models as base learners and utilizing an Extra-Trees meta-learner, SR2P effectively combines complementary predictive patterns across models while maintaining robustness across different spatial and tissue contexts. We show that the prediction performance of the predictive models remains stable in within-sample and within-tissue evaluation settings but decreases markedly in cross-tissue validation. This observation indicates that transcriptome–proteome relationships are strongly tissue-dependent, and models trained on one tissue do not necessarily transfer to others. Moreover, cross-tissue validation results further suggest partial conservation of spatial molecular structure across certain tissue pairs, whereas HNSCC and glioblastoma exhibit markedly reduced performance accompanied by substantially higher variability. This pattern reflects pronounced tissue-specific heterogeneity in both transcriptomic organization and transcriptome–proteome coupling, which limits the transferability of predictive models across anatomically and biologically distinct tissues. Accordingly, training and testing within the same or closely related tissue contexts offer the most reliable strategy for practical protein abundance inference in spatial transcriptomics studies. Spatially augmented models (PLS-Spatial, XGBoost-Spatial, LightGBM-Spatial, and CatBoost-Spatial) outper-form their baseline counterparts (PLS, XGBoost, LightGBM, and CatBoos) across benchmarking settings and tissue types, demonstrating the critical importance of incorporating spatial information into prediction models. Furthermore, GNN-based models generally exhibit lower and more variable predictive performance than non-GNN approaches, suggesting that the existing graph architectures do not yet fully capture the structure-function relationship underlying spatial protein abundance. Regarding speed, the inference time for protein abundance prediction remains low across all methods (on the order of seconds), supporting the practical applicability of SR2P.

The biological application with HNSCC samples from patients treated with immunotherapy demonstrates that predicted protein abundance carries a meaningful biological signal that complements transcriptomic information. Incorporating predicted protein features improves the spatial delineation of macrophage-enriched tumor regions and strengthens the correspondence between inferred cell-type composition and marker gene expression. Moreover, predicted protein profiles distinguish responder and non-responder immune states, with responders exhibiting higher abundance of T cell–associated markers and non-responders enriched for macrophage-associated signatures. These results indicate that predicted protein abundance enables refined characterization of immune microenvironments, even in datasets lacking direct proteomic measurements.

In summary, SR2P provides an efficient framework for spatial protein abundance prediction that integrates spatial information, accommodates heterogeneous modeling strategies, and adapts to tissue-specific transcriptome– proteome relationships. By enabling accurate protein inference and facilitating downstream biological analysis, SR2P expands the analytical capabilities of spatial transcriptomics and offers a practical solution for spatial multi-omics integration when direct protein measurements are unavailable.

## Materials and Methods

### Competing predictive models

We select seven distinct predictive ML methods across three different categories for the comparison, including one linear regression model: partial least squares (PLS) regression ^26^, (2) three tree-based gradient-boosting (ensemble) methods: XGBoost ^27^, LightGBM ^28^, and CatBoost ^29, 30^, and (3) three graph neural network (GNN) methods: GraphSAGE ^32^, GAT ^31^, and DGAT^25^ . Since GNN methods inherently integrate neighborhood information into their models, we apply two settings - spatial and non-spatial-to only the four non-GNN methods. Thus, results of a total of eleven method configurations are reported. We summarize the three categories of predictive ML models below, with further details of each model and its training procedures provided in the Supplementary Methods.

### Linear regression

We use PLS Regression ^26^, which integrates principal component analysis with multiple linear regression. PLS is well suited for high-dimensional omics data due to its ability to handle multi-collinearity effectively ^36^. We build a separate PLS model for each protein for the prediction.

### Tree-based gradient-boosting methods

Gradient boosting models are powerful non-linear predictors that combine multiple decision trees through boosting ^37, 38^ that have been widely adopted in genome-wide analysis in recent years ^39–41^. Three methods including XGBoost ^27^, LightGBM ^28^, and CatBoost ^29, 30^ are selected for evaluation as they are commonly used with computational efficiency and advanced performance in multiple general tasks. All models are trained using the mean squared error (MSE) loss to quantify the discrepancy between predicted and observed protein abundance.

### Graph neural network models

Graph Neural Networks (GNNs) ^42^ extend traditional neural networks to graph-structured data by enabling information exchange across neighboring nodes. This inductive bias makes GNNs particularly suitable for spatial omics analysis, where local tissue architecture and neighborhood context influence molecular phenotypes ^43–45^. We evaluate three GNN models: GAT ^31^, which employs attention-based message passing, and GraphSAGE ^32^, which uses neighborhood aggregation with fixed-size sampling, and Dual-Graph Attention Network (DGAT) ^25^, which integrates both gene co-expression and spatial adjacency graphs to model coordinated regulation. For each tissue section, we construct a spatial graph by connecting each spot to its *k* nearest neighbors based on Euclidean distances between spatial coordinates. All GNN models are trained end-to-end to minimize mean squared error (MSE) in protein abundance prediction.

### Construction of spatial feature for non-GNN models

While GNNs naturally incorporate spatial information through message passing over graphs, traditional prediction models including PLS, XGBoost, LightGBM, and CatBoost do not account for spatial context unless explicitly encoded in the input features. This limitation leads to the loss of valuable spatial information embedded in tissue architecture, ultimately degrading prediction performance on spatial genomics data ^35^.

In this study, we construct spatial features for all non-GNN models (with spatial setting) by incorporating transcriptomic information from each spot’s local neighborhood, constructing four new predictive models: PLS-Spatial, XGBoost-Spatial, LightGBM-Spatial, and CatBoost-Spatial, respectively. Specifically, for each spot, we identify its four neighbors along a fixed order of cardinal directions: west and east neighbors (horizontal axis), and north and south neighbors (vertical axis). These neighbors are identified using exact coordinate offsets from the target spot. If one or more neighbors are missing due to irregular tissue shape or boundary effects, spatially closest unselected neighbors (based on Euclidean distance) are selected instead.

When using the spatial setting for non-GNN models, the RNA expression input of a spot consists of its original gene expression vector concatenated with the vectors from its four selected neighbors, arranged in a fixed spatial order: west, north, east, and south. This fixed ordering preserves the spatial configuration in the expanded feature representation, allowing models to implicitly capture directional spatial dependencies. This strategy enables traditional machine learning models to incorporate spatial structure in a systematic and reproducible way without relying on an explicit graph representation. The same spatial feature construction process is applied uniformly to both the training and testing datasets.

### Stacking framework of SR2P for spatial protein prediction

In SR2P (**Fig. 1a**, and **Suppl Fig. 1**), we implement a stacking-based machine-learning framework ^46, 47^ to integrate multiple predictive models. In bioinformatics and medical applications, stacking has demonstrated advantages in a variety of tasks such as drug response prediction ^48^ and cancer detection ^49^. In SR2P, 11 predictive models described above including PLS, XGBoost, LightGBM, CatBoost, PLS-Spatial, XGBoost-Spatial, LightGBM-Spatial, CatBoost-Spatial, GAT, GraphSAGE and DGAT are used as base learners in the stacking framework. Details of the stacking framework implemented in SR2P are provided in **Suppl Fig. 1**. Briefly, for each base learner, five-fold cross validation is performed on training data to generate out of fold predictions (OOFPs), ensuring that each training sample is predicted by a model that has not seen it during training. The resulting OOFPs from all base learners are collected to form the meta-feature matrix, which is utilized for the input of the meta learner of SR2P. The prediction output of the meta-learner constitutes the final result of SR2P. In SR2P, Extremely randomized trees model (or ExtraTrees) ^50^ is used as the meta-learner, as it demonstrated the best performance among many methods we attempted in our experiments (**Suppl Fig. 16**).

### Cross-validation strategies for evaluating the predictive ML models

We deploy three cross-validation approaches, including within-sample, within-tissue, and cross-tissue validation, on the spatial genomics samples (see section **Datasets**) to assess the accuracy, robustness, and generalizability of the prediction methods.

#### Within-sample validation

Within-sample validation evaluates a model on the same data used for training, providing a measure of fit but sometimes overestimating performance due to the lack of independence between training and test sets. This within-sample cross-validation approach is applied to individual samples. To ensure the preservation of spatial structure during model evaluation, we implement a spatially aware 10-fold cross-validation scheme. Specifically, k-means clustering is applied to the two-dimensional spatial coordinates of all spots of the sample to obtain 10 spatially contiguous groups. This ensures that each fold corresponds to a spatially localized region within the tissue. K-means clustering also preserves approximately equal-sized folds, facilitating balanced evaluation across all cross-validation splits. In each iteration, one spatial cluster is held out as the test set, while the remaining nine clusters serve as the training set.

#### Within-tissue validation

Within-tissue-type validation tests generalizability within the same tissue type by training on one biological sample and testing on another (from the same tissue), capturing variability from technical and sample-specific differences. This approach is applied to the human tonsil and HNSCC datasets (see section **Datasets**). Particularly, for human tonsil datasets, the Tonsil 1 sample is used for training and the Tonsil 2 sample for testing, then the roles are reversed. The same procedure is applied to the HNSCC datasets. This procedure guarantees that all data samples used in the training and evaluation.

#### Cross-tissue validation

Cross-tissue validation challenges further the generalizability of the methods by training and testing across samples from different tissue types, assessing robustness to broader biological diversity, and revealing the model’s ability to handle unseen tissue contexts. This approach is applied to all six spatial genomics samples. In each round, all samples from healthy or diseased tissues are designated as the testing set, and the remaining type of tissues are used for training.

## Marker gene selection from scRNA-seq reference

Marker genes were selected from the scRNA-seq reference data using a two-statistic approach ^35^. For each cell type, we identified genes that are highly up-regulated in the target cell type while showing no significant expression differences among the remaining cell types. Specifically, a robust t-test statistic (T1) was used to test whether the target cell type differs from all other cell types, and a chi-square statistic (T2) was used to assess expression homogeneity across the non-target cell types. Genes were retained as cell-type-specific markers if *p* < 0.05 for T1 and *p* > 0.05 for T2. For each cell type, the top *t* = 10 genes ranked by T1 were selected as marker genes.

### Performance metrics for evaluation

To quantitatively compare model performance across evaluation settings, three complementary metrics are computed for each protein-specific model: Spearman correlation, Pearson correlation, and root mean squared error (RMSE). These metrics are calculated between the predicted and measured protein abundances across all spots in the test set.

Formally, let 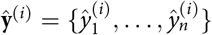 denote the predicted abundance vector for protein *i* from the model *f*^(*i*)^, and 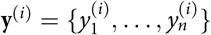 the corresponding measured abundance vector.

#### Spearman correlation

The Spearman rank correlation coefficient 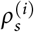 is used as the primary metric to assess the monotonic relationship between ŷ ^(*i*)^ and **y**^(*i*)^. It is computed as:

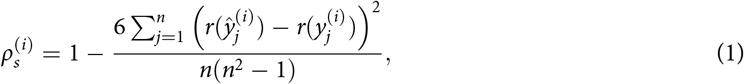

where *r*(·) denotes the rank transformation. Spearman correlation is robust to non-linear dependencies and is reported for all within-sample, within-tissue, and cross-tissue validation experiments.

#### Pearson correlation

The Pearson correlation coefficient 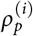 measures the linear association between predic-tions and observations:

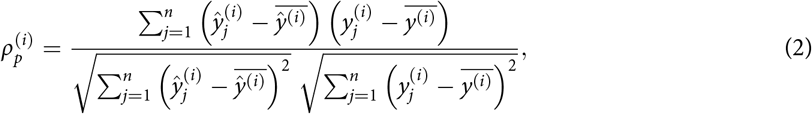

where 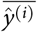 and 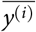 denote the sample means.

#### Root mean squared error (RMSE)

RMSE quantifies the average magnitude of prediction errors:

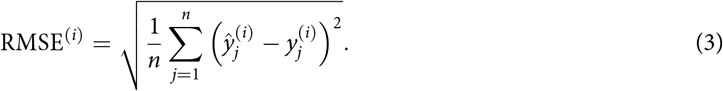

Lower RMSE values indicate more accurate predictions in absolute terms and provide a complementary perspective to correlation-based metrics.

#### Statistical significance testing

To evaluate the effect of incorporating spatial features in non-GNN methods, we compared each base model (non-spatial setting) with the same model in spatial setting using a two-sided Wilcoxon signed-rank test on the per-protein Spearman correlations. For each protein *i*, the paired difference is

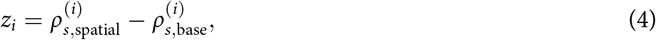

and the Wilcoxon statistic is

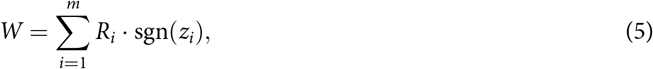

where *R*_*i*_ is the rank of |*z*_*i*_|, *m* is the number of proteins considered, and sgn(·) is a sign function that sgn(*z*_*i*_) = 1 if *z*_*i*_ > 0 and sgn(*z*_*i*_) = −1 if *z*_*i*_ < 0.

## Datasets

### Visium CytAssist multi-omics (RNA+Protein) platform

A total of six spatial genomics samples from the Visium CytAssist multi-omics platform are used in this study. These include four public Tonsil 1, Tonsil 2, breast cancer, and glioblastoma samples obtained from the 10X Genomics webpage ^14^, and two in-house HNSCC samples A1 and D1 (see data generation for these two samples below).

- **Breast cancer**: Formalin-fixed paraffin-embedded (FFPE) human breast cancer tissue (female, 50s) is prepared using the 10x Genomics Visium CytAssist Spatial Gene and Protein Expression technology. Tissue sections of 5 µm thickness are stained with DAPI, Vimentin, and PCNA for immunofluorescence and imaged on an Olympus slide scanning microscope (20× UPLXAPO objective, NA 0.8) with a Hamamatsu Orca FusionBT sCMOS camera. Spatial gene and protein expression libraries are sequenced on an Illumina NovaSeq with targeted depths of 35,000 reads per spot for gene expression and 20,000 reads per spot for protein expression. The dataset comprises 4,169 tissue-covered spots, with a median of 5,473 genes per spot, a mean of 44,009 reads per spot for gene expression, and a mean of 43,038 reads per spot for protein expression.
- **Glioblastoma**: The sample is from a FFPE human glioblastoma tissue (female, 60s) and generated using the same protocol in the breast cancer sample. The dataset comprises 5,756 tissue-covered spots, with a median of 7,629 genes per spot, a mean of 68,906 reads per spot for gene expression, and a mean of 75,377 reads per spot for protein expression.
- **Tonsil 1 & 2**: Both tonsil samples in this dataset are generated using the same protocol above. Tonsil 1 sample is generated from a FFPE human tonsil tissue (female, 30s), comprising 4,194 tissue-covered spots, with a median of 8,275 genes per spot, a mean of 111,046 reads per spot for gene expression, and a mean of 72,898 reads per spot for protein expression. Tonsil 2 sample is based on a FFPE human tonsil tissue of a male with 25 years old. It comprises 4,908 tissue-covered spots, with a median of 8,114 genes per spot, a mean of 75,598 reads per spot for gene expression, and a mean of 34,845 reads per spot for protein expression.
- **Head and neck squamous cell carcinoma (HNSCC)**: Two FFPE human head and neck squamous cell carcinoma tissue samples (A1 and D1) are generated using the 10x Genomics Visium CytAssist Spatial Gene and Protein Expression platform. Tissue sections are prepared following the standard Visium CytAssist FFPE workflow and processed with the Visium Human Transcriptome Probe Kit and the 35-plex Visium Human Cell Profiling Panel. Spatial gene expression and protein libraries are sequenced on an Illumina NovaSeq X Plus platform. Sample A1 comprises 486 tissue-covered spots, with a median of 6 genes per spot, a mean of 169,346 reads per spot for gene expression, and a mean of 130,024 reads per spot for protein expression. Sample D1 comprises 966 tissue-covered spots, with a median of 1,137 genes per spot, a mean of 81,833 reads per spot for gene expression, and a mean of 53,410 reads per spot for protein expression. The dataset has been deposited in GEO and will be publicly available upon acceptance of the manuscript.

### Visium Spatial Gene Expression (only RNA) platform

An independent cohort of 8 HNSCC FFPE tissue samples from patients treated with anti-PD1 ICB ^33^ is included for exploring if the predicted protein markers uncover markers of ICB responses. These samples are profiled using the 10x Genomics Visium Spatial Gene Expression platform and contain only spatial transcriptomic measurements, *without accompanying antibody-derived protein data*. The samples originate from multiple anatomic sites, including the tongue, tonsil, buccal mucosa, oropharynx, tongue base, and cervical lymph node metastasis. Clinical response information to immune checkpoint inhibitor (ICI) therapy is available for all samples, with response status categorized as responder or non-responder (**Suppl Table S2**).

### Data Pre-processing

All six Visium CytAssist datasets provide paired RNA expression and protein abundance of each spot. Low-quality spots are removed if they contain fewer than 700 detected genes or if more than 30% of the unique molecular identifiers (UMIs) are mitochondrial. Genes detected in fewer than 2.5% spots are excluded. RNA counts were library-size normalized to 10,000 total UMIs per spot, log-transformed, scaled to unit variance per gene, and clipped at a maximum value of 10. Protein abundance (antibody-derived tag, ADT counts) is normalized using the centered log-ratio (CLR) transformation recommended by 10x Genomics. After preprocessing, the dataset contains 18,085 genes and 31 proteins across six samples for downstream analyses.

### Prediction evaluation for Visium Spatial Gene Expression (only RNA) data

We apply SR2P, the top-performing method in the within-tissue evaluation to predict protein abundance in eight HNSCC samples generated by the Visium Spatial Gene Expression platform (see section **Datasets**). The model is trained based on the RNA expression and protein abundance produced by Visium CytAssis from both HNSCC samples (A1 and D1). For each sample, we predict protein abundance and construct three feature representations: RNA-only, predicted protein-only, and combination which integrates RNA expression with predicted protein abundance. We perform spatial clustering separately for each representation using the DECLUST pipeline 35 to derive spatially coherent tissue domains. Following clustering, we apply the DECLUST deconvolution to each cluster to estimate cluster-level cell-type compositions, including macrophage proportions. In parallel, we compute the mean expression of macrophage-associated RNA markers (e.g., *CD68, CD163, MSR1*) at the cluster level. We then evaluate the correspondence between cluster-level macrophage proportions and marker gene expression to assess whether the inclusion of predicted protein abundance improves the characterization of the tumor immune microenvironment. Cluster-level comparisons across samples are assessed using two-tailed Wilcoxon rank-sum tests.

## Code availability

The SR2P software is available at https://github.com/Qingyueee/SR2P

## Data availability

This study utilized publicly available datasets and newly generated in-house data, which will be made available upon publication.

The publicly available datasets, including the:

Dataset1 (Human Breast Cancer): 10X Visium CytAssist, Human-Breast-Cancer).

Dataset2 (Human Glioblastoma): 10X Visium CytAssist, Human-Glioblastoma).

Dataset3 (Human Tonsil 1): 10X Visium CytAssist, Human-Tonsil-1).

Dataset4 (Human Tonsil 2): 10X Visium CytAssist, Human-Tonsil-2).

## Acknowledgements

This work was partially supported by funding from the Swedish Research Council (VR) and CancerFonden [22 2020 Pj & 25 4528 Pj]. Research reported in this publication was partially supported by the Wisconsin Head & Neck Cancer SPORE (P50CA278595) Career Enhancement Program pilot grant to H.Q.D. This research was also supported by grants from the National Institutes of Health (R35GM150893, P01CA022443 to H.Q.D). We thank Dr Darya Buehler at the Department of Pathology and Laboratory Medicine, UW-Madison School of Medicine and Public Health, for advice in selecting tissue-captured areas. The computations were performed using resources provided by the National Academic Infrastructure for Supercomputing in Sweden (NAISS), which is partially funded by the Swedish Research Council through grant agreement no. 2025-22-991 and 2025-5-655. Qingyue Wang is grateful for the financial support from the China Scholarship Council (CSC).

## Author contributions

H.Q.D and T.N.V conceptualized the study. Q.W was responsible for algorithm development and implementation with inputs from T.N.V and Y.P. Analysis was conducted by Q.W, Y.L with contributions from P.K, T.N.V and H.Q.D. In-house Visium CITE-Seq were generated by A.G, R.H. The initial draft of the manuscript was written by Q.W, H.Q.D and T.N.V with reviewing and editing from all other authors. Supervision was provided by T.N.V, H.Q.D, J.H and Y.P.

## Ethics declarations

Utilization of human tissue in this study was approved by University of Wisconsin Institutional Review Board (IRB approval number 2018-1510)

