## Supplemental material for "SR2P: an efficient stacking method to predict protein abundance from gene expression in spatial transcriptomics data"

### Supplementary Methods

#### Tissue preparation and spatial gene and protein profiling

Sections with 10 μm thickness for de-identified head and neck cancer patient tissue samples from the Wisconsin Head and Neck Cancer SPORE were placed on individual positively charged slides. The tissues were sectioned following the 10x Genomics Visium CytAssist Spatial Gene and Protein Expression FFPE Tissue Preparation Guide (Demonstrated Protocol CG000660, Rev A). Slides with 6.5 mm × 6.5 mm capture area cassettes were used, and tissue capture areas were selected. Sections were deparaffinized, stained with hematoxylin and eosin, imaged on an Aperio scanner at 40× magnification, decoverslipped, destained, and decrosslinked according to the protocol described (CG000658, Rev A).

Following DNase treatment, samples underwent probe hybridization with the Visium Human Transcriptome Probe Kit, probe ligation, and protein labeling with the 35-plex Visium Human Cell Profiling Panel using the 10x Genomics Visium CytAssist Spatial Gene and Protein Expression for FFPE workflow (Demonstrated Protocol CG000494, Rev C). Tissue slides were then loaded into a CytAssist Instrument, where gene expression probes and antibody tags were released together with spatial barcode information following probe and tag extension. Paired gene expression and protein libraries were generated outside the instrument following the standard Visium FFPE workflow and sequenced with paired-end dual indexing on an Illumina NovaSeq X Plus platform (PE150).

Manual fiducial alignment and tissue outlining were performed in Loupe Browser to ensure accurate detection of tissue boundaries. The aligned images were exported and processed using the Space Ranger pipeline (v2022.0705.1, 10x Genomics) with alignment to the GRCh38-2020-A reference genome. Sample A1 comprises 486 tissue-covered spots, with a median of 6 genes per spot, a mean of 169,346 reads per spot for gene expression, and a mean of 130,024 reads per spot for protein expression. Sample D1 comprises 966 tissue-covered spots, with a median of 1,137 genes per spot, a mean of 81,833 reads per spot for gene expression, and a mean of 53,410 reads per spot for protein expression.

#### Linear Regression Baseline

**PLS Regression.** PLS models the relationship between the RNA expression matrix $\mathbf{X}$ ($n \times g$) and each protein abundance vector $\mathbf{Y}^{(i)}$ by projecting both predictors and responses into a shared latent space:

$$\mathbf{X} = \mathbf{T}\mathbf{P}^\top + \mathbf{E}, \quad \mathbf{Y}^{(i)} = \mathbf{T}\mathbf{q}^\top + \mathbf{F},$$

where $\mathbf{T}$ contains latent components that maximize the covariance between $\mathbf{Xw}$ and $\mathbf{Y}^{(i)}$, subject to $\|\mathbf{w}\| = 1$. The regression coefficients are computed as:

$$\widehat{\beta}_{\text{PLS}} = \mathbf{W}(\mathbf{P}^\top\mathbf{W})^{-1}\mathbf{q}, \quad \text{and} \quad \widehat{\mathbf{Y}}^{(i)} = \mathbf{X}\widehat{\beta}_{\text{PLS}}.$$

A separate PLS model was trained for each protein using 5 latent components.

#### Tree-Based Ensemble Models

All tree-based ensemble boosting models used in this study follow a shared formulation:

$$\hat{y}_j^{(i)} = \sum_{k=1}^{K} f_k(\mathbf{x}_j), \quad f_k \in \mathcal{F},$$

where $f_k$ are regression trees, and $\mathbf{x}_j$ is the input for spot $j$. Model optimization minimizes the regularized squared error:

$$\mathcal{L}^{(i)} = \sum_{j=1}^{n} (y_j^{(i)} - \hat{y}_j^{(i)})^2 + \sum_{k=1}^{K} \Omega(f_k), \quad \Omega(f_k) = \gamma T_k + \frac{1}{2}\lambda\|w_k\|^2.$$

All boosting models were trained separately for each protein with default hyperparameters, using a fixed random seed and full parallelism enabled.

**XGBoost.** XGBoost employs a second-order Taylor expansion of the loss and sparse-aware split finding. The objective at each iteration is approximated as:

$$\mathcal{L}^{(t)} \approx \sum_{j=1}^{n} \left[ g_j f_t(\mathbf{x}_j) + \frac{1}{2} h_j f_t^2(\mathbf{x}_j) \right] + \Omega(f_t),$$

where $g_j$ and $h_j$ are the first and second derivatives of the loss function. The model was trained using the squared error loss.

**LightGBM.** LightGBM constructs trees using a leaf-wise growth strategy that selects splits with the largest loss reduction. It employs histogram-based feature binning and gradient-based one-side sampling (GOSS) to reduce training complexity and accelerate convergence. Default parameters were used with squared error loss.

**CatBoost.** CatBoost builds symmetric trees and uses permutation-driven ordered boosting to avoid overfitting and target leakage. While originally designed for categorical features, it also performs competitively on fully numerical input. The model was trained with root mean squared error (RMSE) as the objective and default parameter settings.

#### Graph Neural Network Models

Graph neural network Models were trained to learn spatial context by constructing a graph $\mathcal{G} = (\mathcal{V}, \mathcal{E})$ for each sample, where nodes $\mathcal{V}$ represent spatial spots, and edges $\mathcal{E}$ are defined via $k$-nearest neighbors in Euclidean space. Node features were derived from gene expression vectors. GNNs were trained separately for each protein using stochastic gradient descent to minimize mean squared error, with a learning rate of 0.001.

**GAT.** GAT updates node features via attention-weighted aggregation:

$$\mathbf{h}'_v = \sigma \left( \sum_{u \in \mathcal{N}(v)} \alpha_{vu} \mathbf{W} \mathbf{h}_u \right),$$

where the attention weights are given by:

$$\alpha_{vu} = \frac{\exp\left(\text{LeakyReLU}(\mathbf{a}^\top [\mathbf{W}\mathbf{h}_v \| \mathbf{W}\mathbf{h}_u])\right)}{\sum_{k \in \mathcal{N}(v)} \exp\left(\text{LeakyReLU}(\mathbf{a}^\top [\mathbf{W}\mathbf{h}_v \| \mathbf{W}\mathbf{h}_k])\right)}.$$

The model employed two attention heads per layer, ELU activation, and dropout between layers.

**GraphSAGE.** GraphSAGE updates node embeddings through sampling-based aggregation:

$$\mathbf{h}'_v = \sigma \left( \mathbf{W} \cdot \text{AGGREGATE} \left( \{\mathbf{h}_v\} \cup \{\mathbf{h}_u : u \in \mathcal{N}(v)\} \right) \right),$$

where AGGREGATE is a mean function, $\sigma$ is ReLU, and $\mathbf{W}$ is a learnable weight matrix. Two aggregation layers were used, followed by a linear regression head.

**DGAT.** DGAT that integrates two heterogeneous graphs—a spatial graph $\mathcal{G}_s$ and an expression graph $\mathcal{G}_e$—to jointly capture spatial proximity and transcriptomic similarity. For each spot $v$, the embeddings from the two graphs are updated by separate GAT layers:

$$\mathbf{h}_v^{(s)} = \mathrm{GAT}_s(\mathbf{h}_v, \mathcal{N}_s(v)), \quad \mathbf{h}_v^{(e)} = \mathrm{GAT}_e(\mathbf{h}_v, \mathcal{N}_e(v)),$$

where $\mathcal{N}_s(v)$ and $\mathcal{N}_e(v)$ denote the neighbors of $v$ in the spatial and expression graphs, respectively.

The two embeddings are combined via a learnable gating parameter $g \in (0, 1)$:

$$\mathbf{h}'_v = g \cdot \mathbf{h}_v^{(s)} + (1 - g) \cdot \mathbf{h}_v^{(e)}.$$

The final latent embedding $\mathbf{z}_v$ is fed into two decoder branches: a protein prediction head and an RNA reconstruction head:

$$\hat{\mathbf{y}}_v^{(\mathrm{prot})} = f_{\mathrm{prot}}(\mathbf{z}_v), \qquad \hat{\mathbf{x}}_v^{(\mathrm{rna})} = f_{\mathrm{rna}}(\mathbf{z}_v).$$

where $f_{\mathrm{prot}}$ is a regression network outputting protein abundances, and $f_{\mathrm{rna}}$ reconstructs the original gene expression vector to regularize the latent space.

The overall training objective is a weighted combination of protein prediction loss and RNA reconstruction loss:

$$\mathcal{L} = \mathcal{L}_{\mathrm{prot}} + \lambda_{\mathrm{recon}} \mathcal{L}_{\mathrm{rna}}.$$

with $\mathcal{L}_{\mathrm{prot}} = \mathrm{MSE}(\hat{\mathbf{y}}^{(\mathrm{prot})}, \mathbf{y}^{(\mathrm{prot})})$ and $\mathcal{L}_{\mathrm{rna}} = \mathrm{MSE}(\hat{\mathbf{x}}^{(\mathrm{rna})}, \mathbf{x}^{(\mathrm{rna})})$.

### Supplementary Tables

**Table S1. Pairwise comparison results between responders and non-responders at the cluster level.**

| Protein | p-value | Adjusted p-value (FDR) |
|---|---|---|
| PTPRC2 | $4.759552 \times 10^{-32}$ | $1.475461 \times 10^{-30}$ |
| CXCR5 | $3.481524 \times 10^{-23}$ | $5.396362 \times 10^{-22}$ |
| CD274 | $4.339586 \times 10^{-21}$ | $3.363179 \times 10^{-20}$ |
| MS4A1 | $2.739636 \times 10^{-14}$ | $1.213267 \times 10^{-13}$ |
| PTPRC | $1.586430 \times 10^{-14}$ | $8.196557 \times 10^{-14}$ |
| PAX5 | $7.358431 \times 10^{-16}$ | $4.562227 \times 10^{-15}$ |
| CD27 | $6.657509 \times 10^{-11}$ | $2.579785 \times 10^{-10}$ |
| ITGAM | $3.314074 \times 10^{-4}$ | $7.338306 \times 10^{-4}$ |
| CD8A | $3.736547 \times 10^{-8}$ | $1.287033 \times 10^{-7}$ |
| CD19 | $3.227955 \times 10^{-4}$ | $7.338306 \times 10^{-4}$ |
| CEACAM8 | $4.164777 \times 10^{-4}$ | $8.607206 \times 10^{-4}$ |
| CCR7 | $4.966172 \times 10^{-1}$ | $5.498262 \times 10^{-1}$ |
| CD4 | $1.664853 \times 10^{-2}$ | $2.580521 \times 10^{-2}$ |
| KRT5 | $1.527228 \times 10^{-4}$ | $4.304007 \times 10^{-4}$ |
| ACTA2 | $5.182900 \times 10^{-4}$ | $1.004187 \times 10^{-3}$ |
| CR2 | $8.069796 \times 10^{-1}$ | $8.338789 \times 10^{-1}$ |
| CD3E | $5.873657 \times 10^{-1}$ | $6.278737 \times 10^{-1}$ |
| HLA-DRA | $1.209455 \times 10^{-1}$ | $1.630135 \times 10^{-1}$ |
| CD40 | $3.890557 \times 10^{-2}$ | $5.743204 \times 10^{-2}$ |
| PCNA | $1.410351 \times 10^{-1}$ | $1.821703 \times 10^{-1}$ |
| BCL2 | $1.198689 \times 10^{-7}$ | $3.715937 \times 10^{-7}$ |
| EPCAM | $3.015942 \times 10^{-1}$ | $3.595931 \times 10^{-1}$ |
| SDC1 | $8.537664 \times 10^{-1}$ | $8.537664 \times 10^{-1}$ |
| PECAM1 | $1.544639 \times 10^{-1}$ | $1.915352 \times 10^{-1}$ |
| CD163 | $4.238029 \times 10^{-1}$ | $4.865886 \times 10^{-1}$ |
| VIM | $1.019148 \times 10^{-2}$ | $1.662820 \times 10^{-2}$ |
| FCGR3A | $2.576045 \times 10^{-4}$ | $6.654784 \times 10^{-4}$ |
| ITGAX | $2.924025 \times 10^{-3}$ | $5.035821 \times 10^{-3}$ |
| CD14 | $1.325004 \times 10^{-3}$ | $2.416183 \times 10^{-3}$ |
| PDCD1 | $6.902013 \times 10^{-2}$ | $9.725564 \times 10^{-2}$ |
| CD68 | $2.215664 \times 10^{-21}$ | $2.289519 \times 10^{-20}$ |

**Table S2. Clinical information for the HNSC spatial transcriptomics validation cohort.**

| **Sample ID** | Age | Gender | **Anatomic Site** | **Immunotherapy Response** |
|---|---|---|---|---|
| ck17_5 | 61 | M | Tonsil | Y |
| ck17_7 | 47 | M | Tonsil | N |
| ck17_12e6 | 68 | M | Tongue | N |
| ck17_19 | 54 | M | Oropharynx | N |
| ck17_25 | 56 | M | Tongue | Y |
| ck17_208 | 49 | M | Buccal | N |
| ck17_1294 | 70 | M | Tongue Base | Y |
| ck17_1592 | 54 | M | Tongue | Y |
| A1 | 68 | M | larynx | Y |
| D1 | 71 | M | hypopharynx | N |

### Supplementary Figures

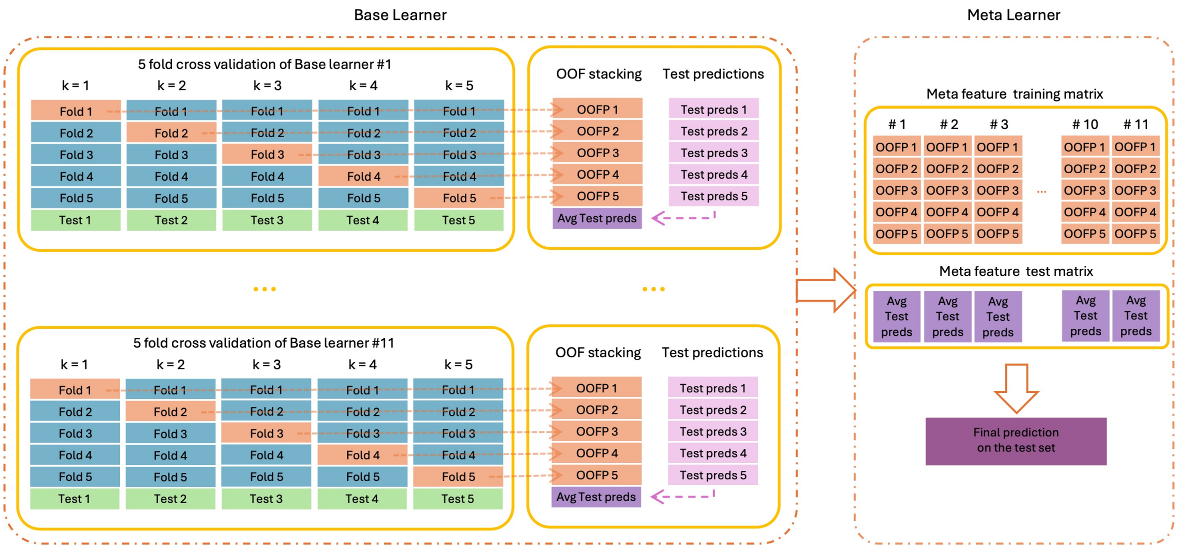


**Figure S1. Overview of the stacking framework used in SR2P.** Schematic illustration of the stacking strategy adopted in this study. For each base learner, five fold cross validation is performed on the training data to generate out of fold predictions (OOFPs), ensuring that each training sample is predicted by a model that has not seen it during training. The resulting OOFPs from all base learners are concatenated to form the meta feature matrix, which is used to train the meta learner. In parallel, each base learner produces predictions on the test set across all folds, which are averaged to obtain stable test predictions. The final prediction on the test set is then generated by applying the trained meta learner to the aggregated test predictions.

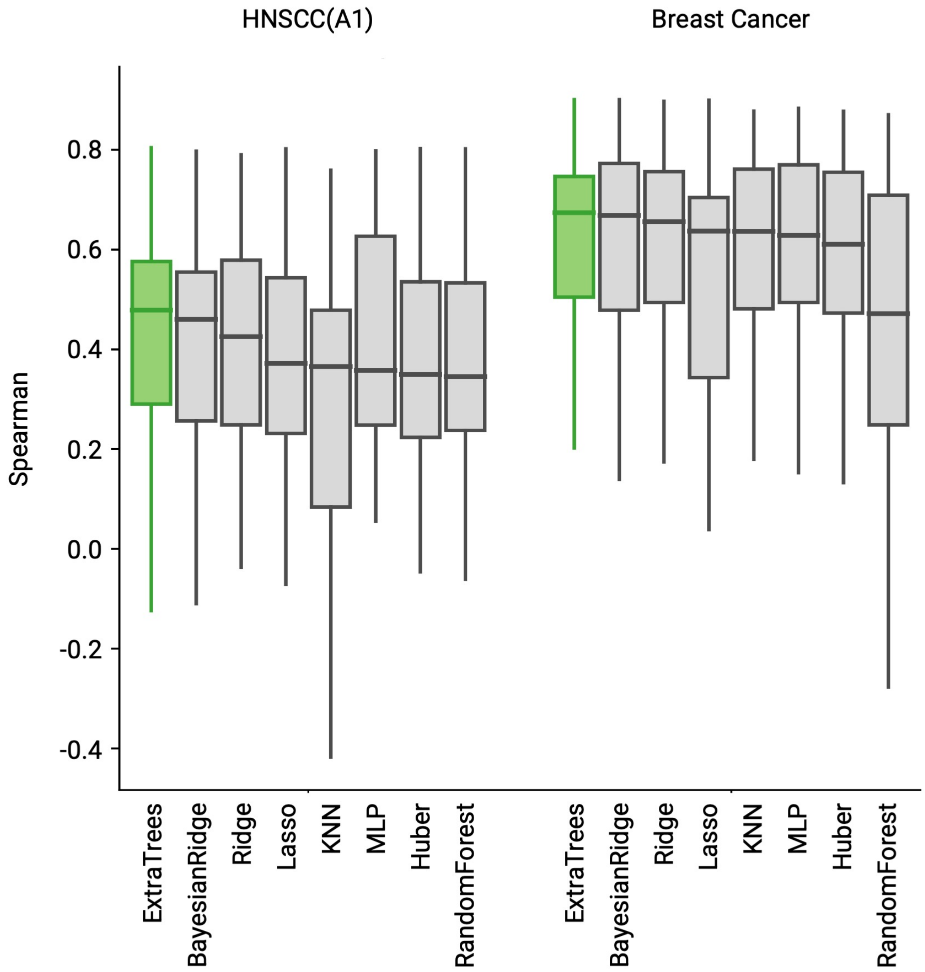


**Figure S2. Performance comparison of candidate meta learners in the stacking framework of SR2P.** Boxplots show the distribution of Spearman correlation between predicted and measured protein expression across proteins in two representative datasets (HNSCC(A1) and Breast Cancer), using different models as the meta learner to integrate out of fold predictions from all base learners. ExtraTrees achieved the best overall performance among the tested meta learners (BayesianRidge, Ridge, Lasso, KNN, MLP, Huber, and RandomForest), and was therefore selected as the final meta learner in SR2P.

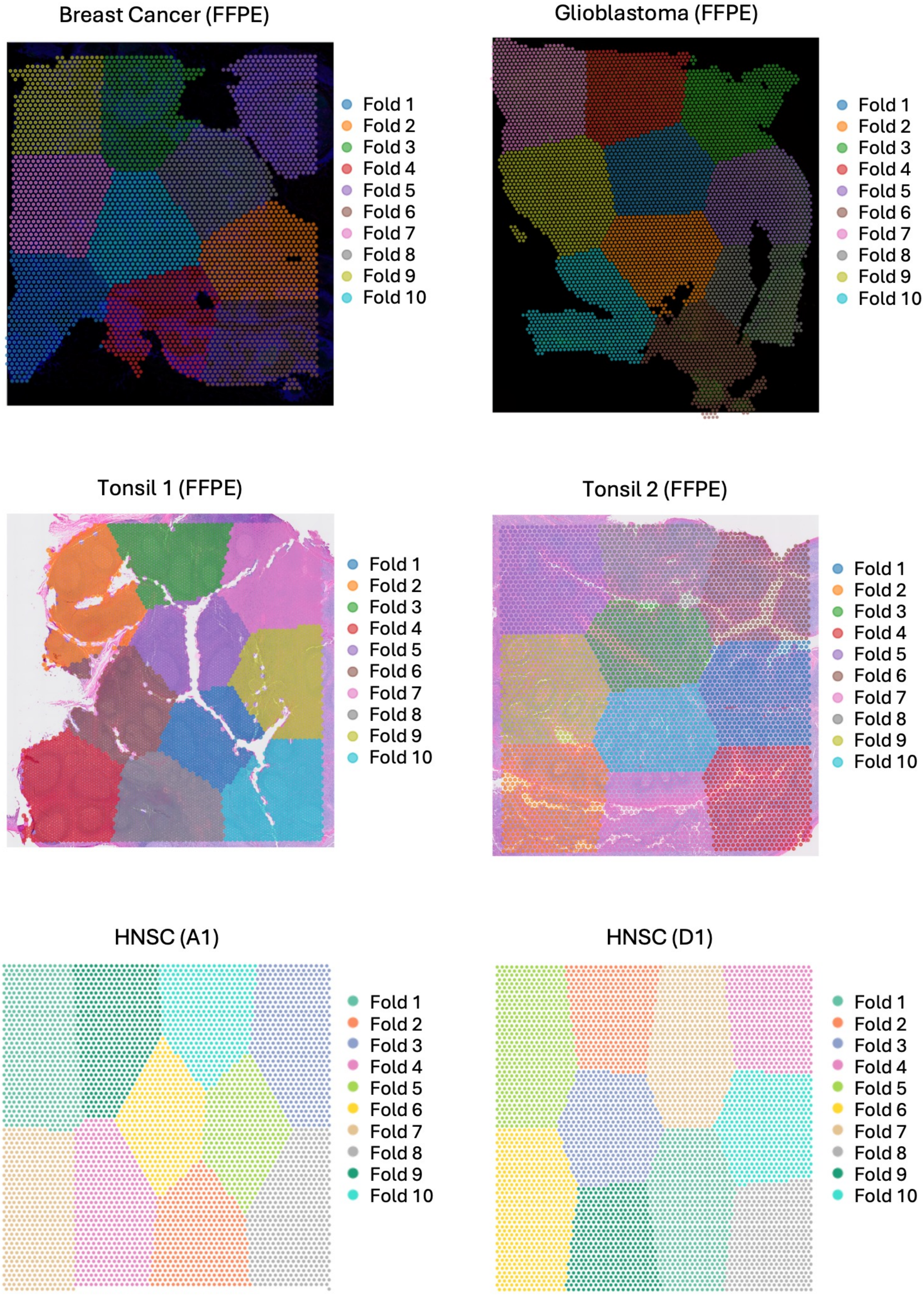
Breast Cancer (FFPE)
Fold 1
Fold 2
Fold 3
Fold 4
Fold 5
Fold 6
Fold 7
Fold 8
Fold 9
Fold 10
Glioblastoma (FFPE)
Fold 1
Fold 2
Fold 3
Fold 4
Fold 5
Fold 6
Fold 7
Fold 8
Fold 9
Fold 10
Tonsil 1 (FFPE)
Fold 1
Fold 2
Fold 3
Fold 4
Fold 5
Fold 6
Fold 7
Fold 8
Fold 9
Fold 10
Tonsil 2 (FFPE)
Fold 1
Fold 2
Fold 3
Fold 4
Fold 5
Fold 6
Fold 7
Fold 8
Fold 9
Fold 10
HNSC (A1)
Fold 1
Fold 2
Fold 3
Fold 4
Fold 5
Fold 6
Fold 7
Fold 8
Fold 9
Fold 10
HNSC (D1)
Fold 1
Fold 2
Fold 3
Fold 4
Fold 5
Fold 6
Fold 7
Fold 8
Fold 9
Fold 10

**Figure S3. Spatial 10-fold cross-validation splits for within-sample evaluation.** Illustration of the spatial partitioning strategy applied to spatial multi-omics datasets for within-sample evaluation. Capture spots were divided into ten spatially contiguous folds using K-means clustering on two-dimensional spatial coordinates, ensuring spatial separation between training and test sets in each fold. Colors represent fold assignments, with each fold serving once as the test set. This spatially aware design preserves tissue architecture and provides a more realistic estimate of model generalization compared to random splitting.

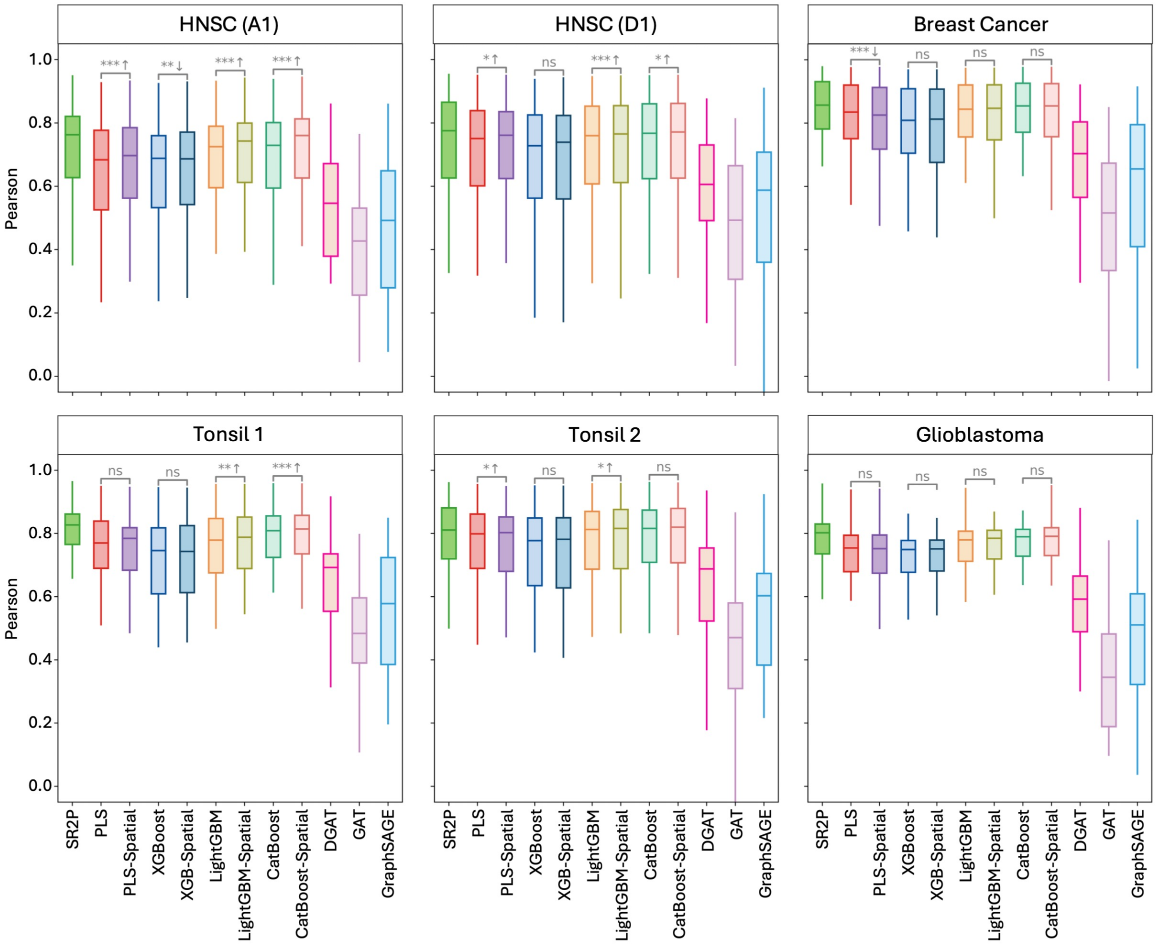


**Figure S4. Prediction accuracy across models (Pearson correlation).** Prediction accuracy of eleven models for each dataset, evaluated using Pearson correlation between predicted and measured protein abundances. The two-tailed Wilcoxon rank-sum test is used to assess differences between spatial and non-spatial settings for non-GNN methods ($p < 0.05$ (*), $p < 0.01$ (**), and $p < 0.001$ (***)), with *ns* indicating no significant difference). Arrows indicate the direction of performance change, where upward (downward) arrows denote that the spatial setting performs better (worse) than the non-spatial setting. SR2P consistently achieves the highest predictive performance across all datasets and models.

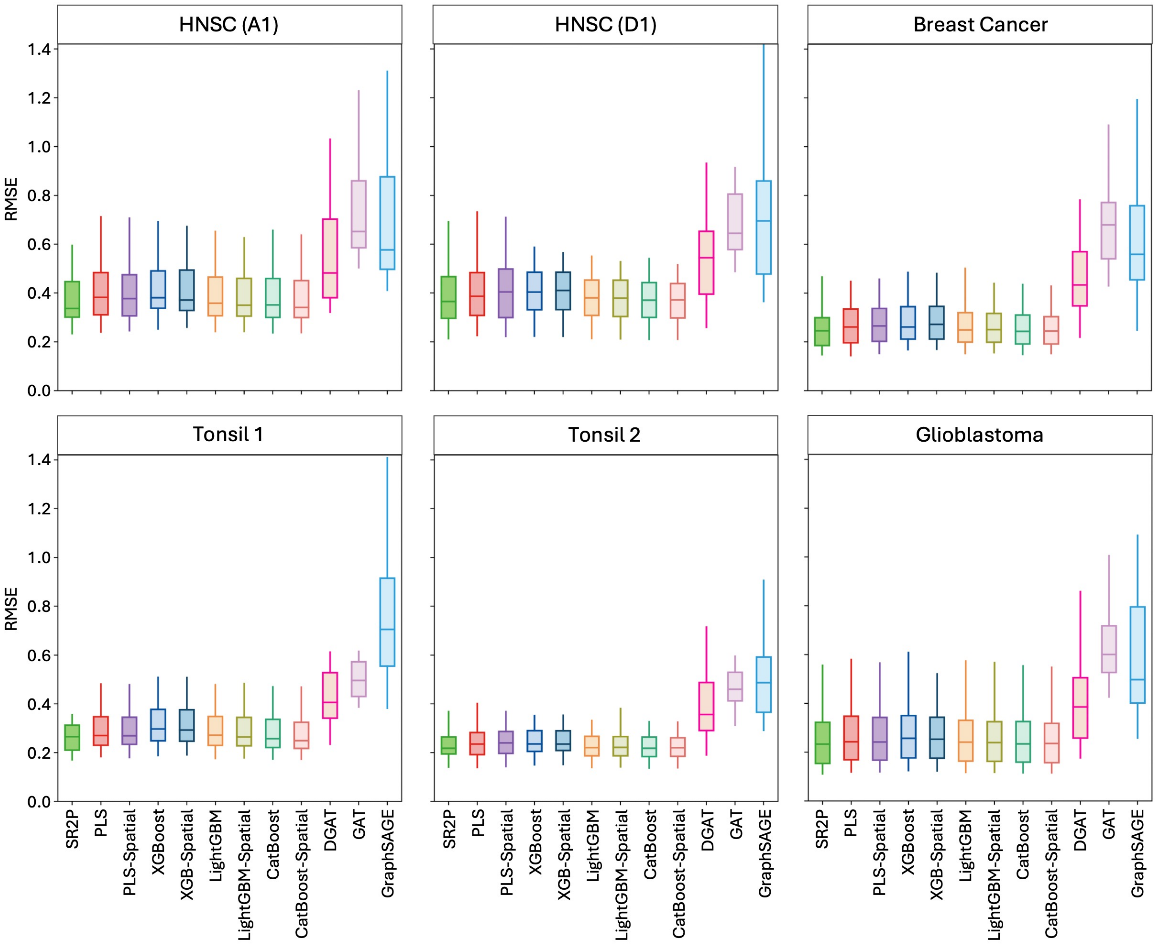


**Figure S5. Prediction error across models (RMSE).** Prediction error of eleven models for each dataset, evaluated using root mean squared error (RMSE) between predicted and measured protein abundances. Lower RMSE values indicate better prediction performance. Across all datasets, SR2P consistently achieves the lowest prediction error, while GNN based methods generally exhibit higher RMSE and greater variability.

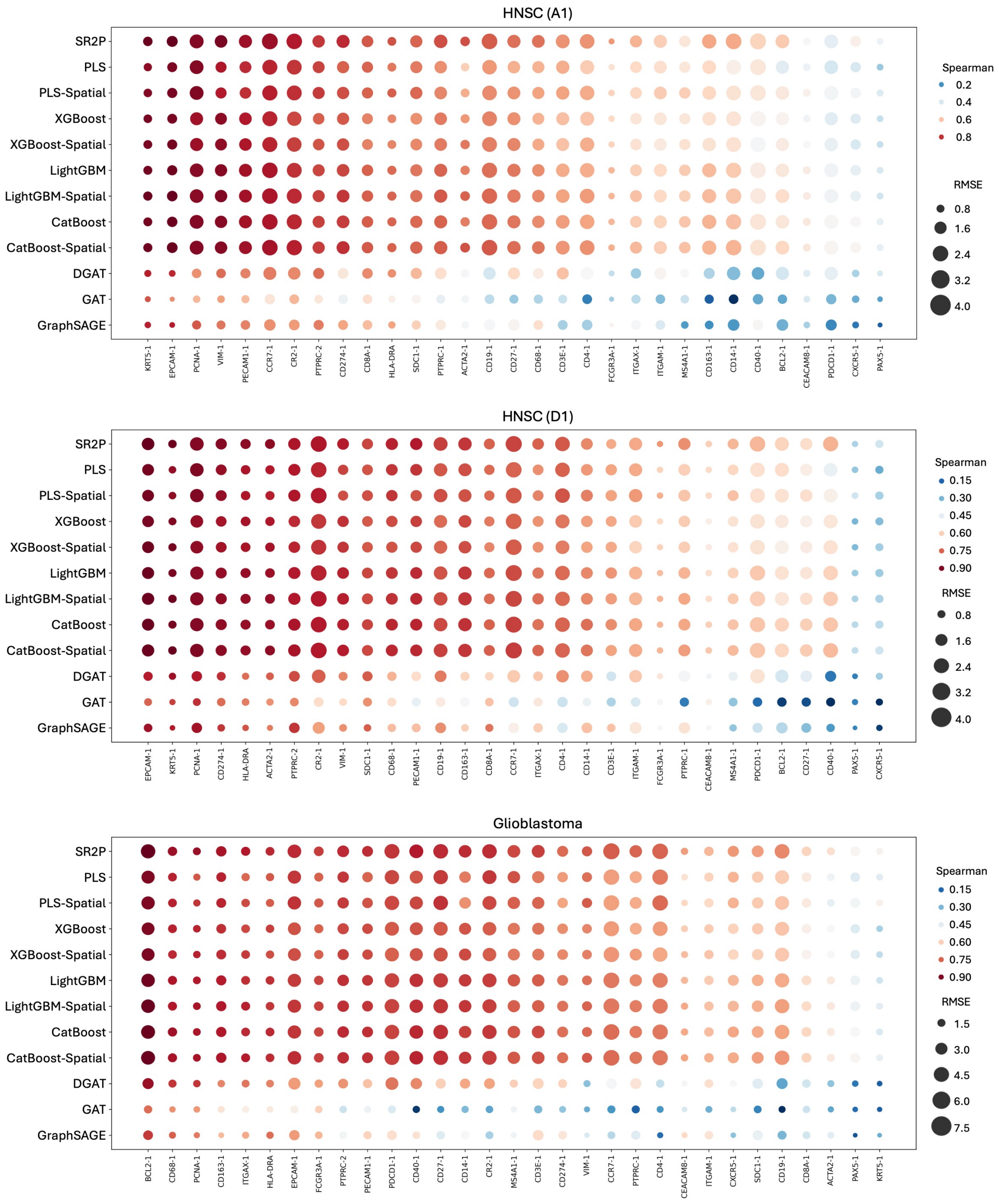


**Figure S6. Protein-wise prediction performance in tumor tissues (Head and Neck Cancer and Glioblastoma).** The figure compares the prediction performance of multiple models across tumor samples. In the HNSC A1 and HNSC D1, proteins associated with epithelial identity and proliferative activity, including EPCAM, KRT5, PCNA, and VIM, show relatively high prediction accuracy, particularly when spatial information is incorporated into the modeling framework. In the glioblastoma sample, proteins reflecting inflammatory and microglial activation states, such as BCL2, CD68, PCNA, and CD163, exhibit stronger predictive performance, consistent with the dominant immune–stromal characteristics of the glioblastoma microenvironment. Notably, PAX5 shows consistently poor prediction performance across all three samples, reflecting either weak transcript–protein coupling or limited spatial predictability for this marker. Overall, models incorporating spatial neighborhood structure demonstrate improved robustness in predicting protein expression in heterogeneous tumor tissues.

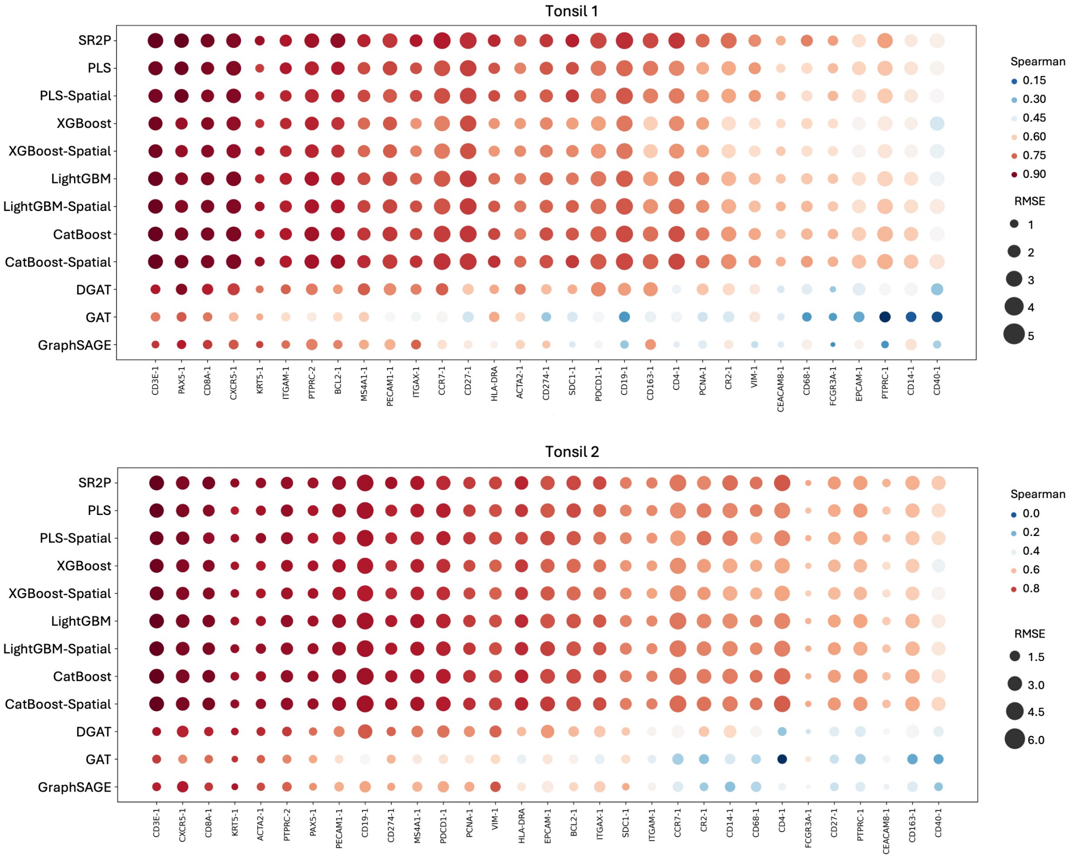


**Figure S7. Protein-wise prediction performance in healthy tonsil tissue.** The figure compares the prediction performance of different models across two healthy tonsil samples. Each panel shows protein-level prediction accuracy, where color represents the Spearman correlation between predicted and measured expression, and point size corresponds to the RMSE. Proteins associated with T cell populations, including CD3E, CD8A, and the pan-leukocyte marker PTPRC2, exhibit consistently high prediction accuracy across both samples, reflecting the well-organized T cell zones in tonsil tissue. In contrast, proteins such as CEACAM8 and FCGR3A show reduced and more variable prediction performance, suggesting weaker transcript–protein coupling or greater spatial heterogeneity for these markers. Overall, the healthy tonsil samples demonstrate robust and reproducible prediction performance for proteins that define canonical immune microanatomical structures.

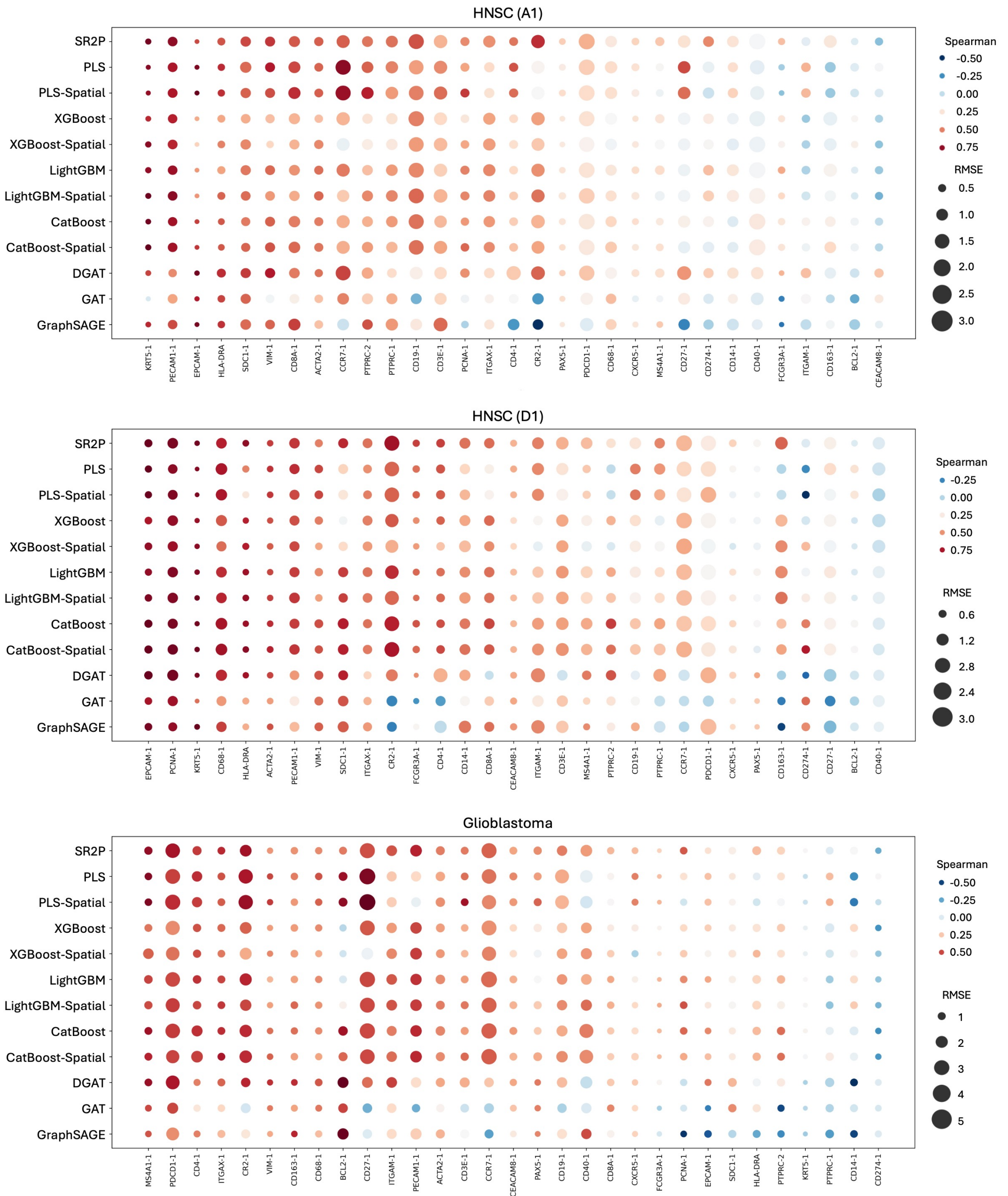


**Figure S8. Cross-tissue protein prediction performance in tumor samples.** Each panel summarizes protein-level cross-tissue prediction performance for HNSC A1, HNSC D1, and one glioblastoma sample. Rows correspond to prediction models and columns to proteins. In the HNSC samples, the epithelial and proliferation-associated markers EPCAM and KRT5 retain comparatively stronger predictive signals, whereas proteins such as CD163 and BLC2 show consistently weak performance. In the glioblastoma sample, macrophage and microglial activation markers including MS4A1, PDCD1, and CD4 maintain relatively higher correlations, while proteins such as CD274 and CD14 display poor transferability across tissues.

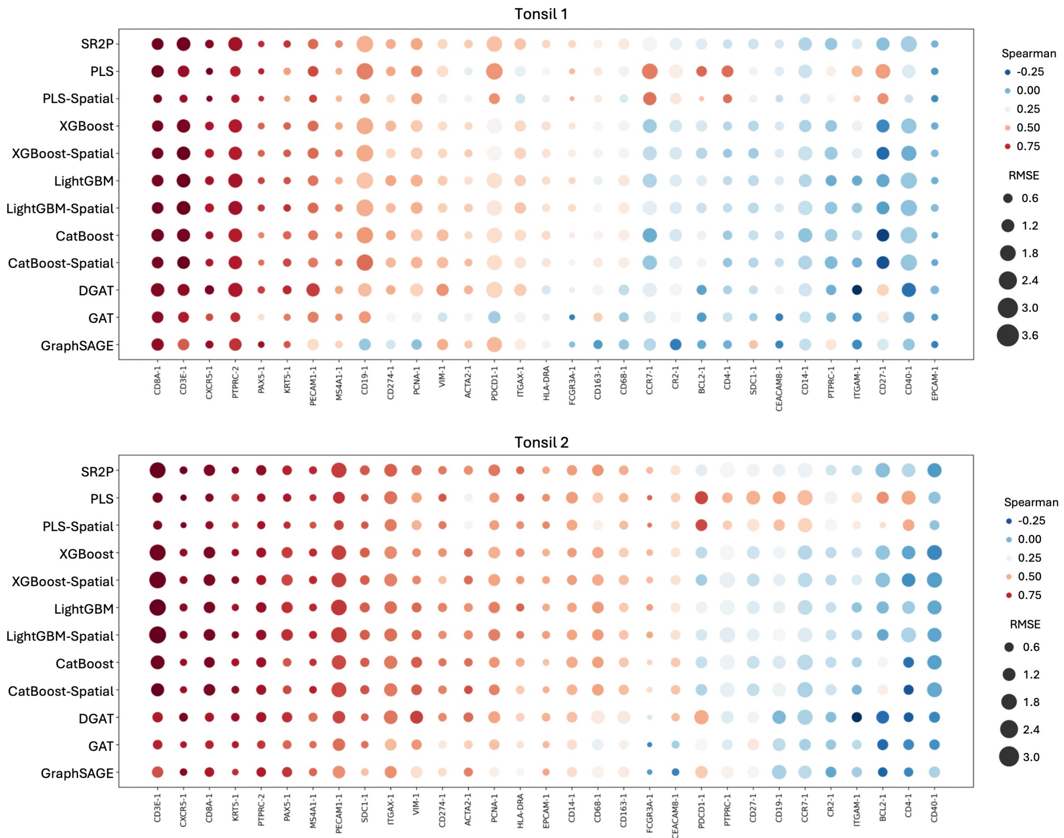


**Figure S9. Cross-tissue protein prediction performance in healthy tonsil samples.** Each panel summarizes protein-level cross-tissue prediction performance for two healthy tonsil samples. Rows correspond to prediction models and columns to proteins. In both samples, the T cell–associated markers CD8A, CD3E, CXCR5, and PTPRC2 retain comparatively stronger predictive signals, whereas proteins such as CD40 and ITGAM exhibit consistently weak performance across models and do not transfer well across tissue contexts.

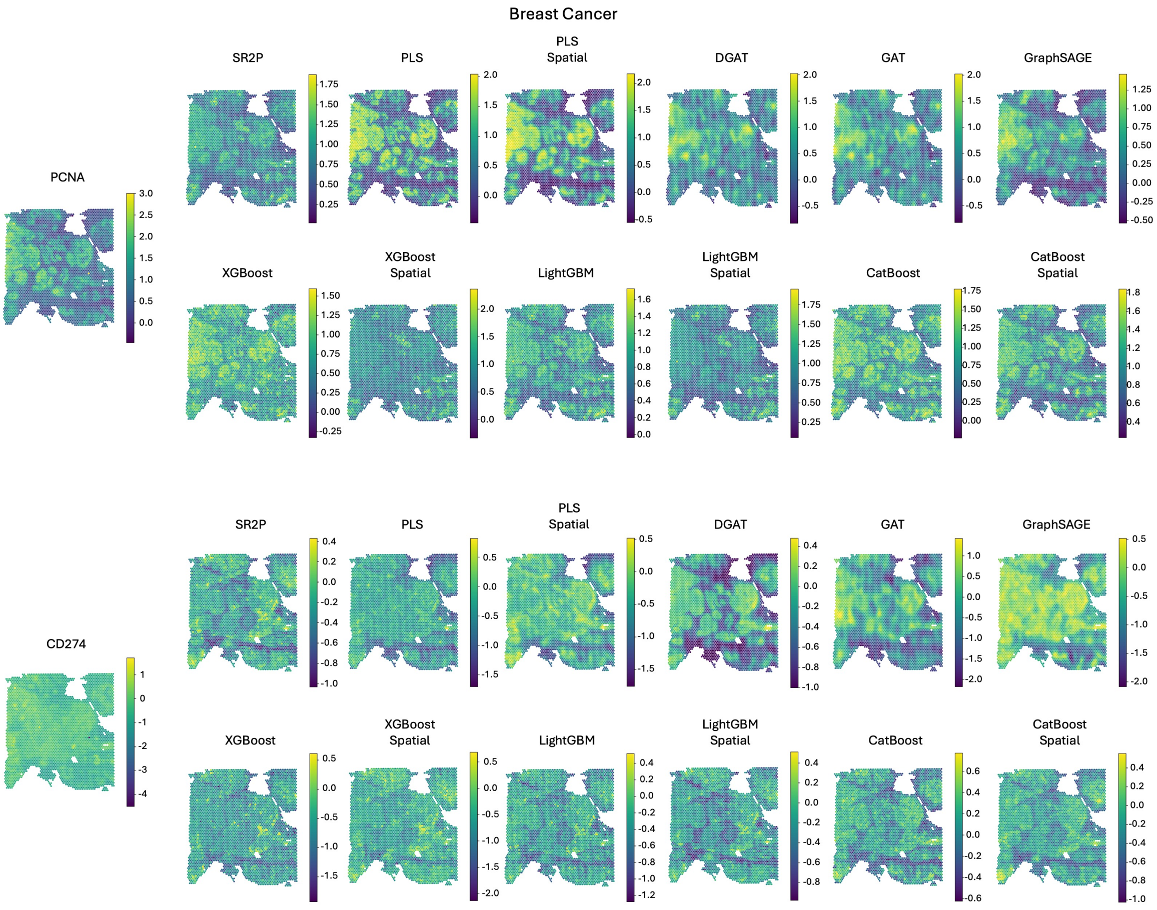


**Figure S10. Cross-tissue prediction of PCNA and CD274 expression in breast cancer.** PCNA is a nuclear marker associated with cell proliferation and exhibits discrete high expression clusters in the measured data. SR2P reproduces the spatial arrangement of these clusters and preserves the overall contrast between high and low expression regions, although peak intensities are slightly compressed. PLS and PLS Spatial generate artificially regular high expression patterns and expand localized signals into broad regions. GNN based methods (DGAT, GAT, and GraphSAGE) smooth out the discrete clusters and fail to recover clear cluster boundaries. Tree based models partially capture block like structures, with CatBoost and its spatial variant showing closer agreement, but local intensities and boundaries remain inconsistent with the measured map. CD274 (PD L1) shows a more diffuse spatial distribution with limited local variation. SR2P maintains the weak spatial gradients observed in the measured data without introducing spurious high or low expression regions. In contrast, PLS based models introduce granular noise or upward intensity shifts, DGAT produces large low expression areas not present in the reference, and GAT and GraphSAGE generate spatially inconsistent patterns. Tree based models yield more homogeneous predictions, often attenuating the subtle spatial variation of CD274.

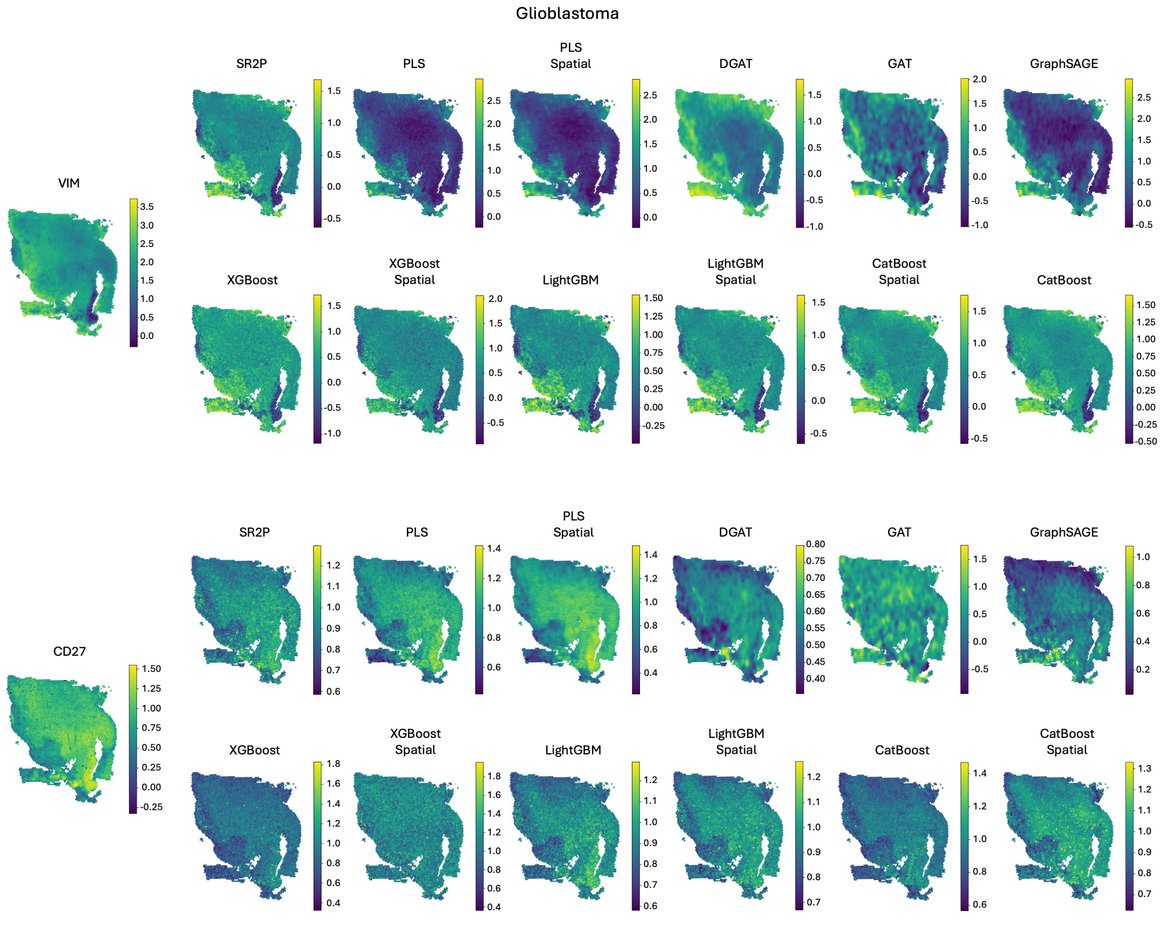


**Figure S11. Cross-tissue prediction of VIM and CD27 expression in glioblastoma.** VIM (vimentin) is a cytoskeletal marker associated with mesenchymal phenotype and tumor invasiveness. SR2P matches the measured map in both the global intensity layout and the location of high expression regions, including the high expression band in the lower part of the tissue and the lower expression area on the right side. PLS and GraphSAGE under predict VIM overall and compress spatial contrast, producing maps that are dominated by low values. GAT introduces strong spatial artifacts and breaks the measured smooth structure into patchy patterns. DGAT captures the coarse gradient but misses part of the low expression region and reduces boundary sharpness. XGBoost, LightGBM, and CatBoost produce visually reasonable reconstructions, and the spatial variants mainly help recover local contrast but do not fully match the reference in all regions. CD27 is a marker of activated T cells. SR2P preserves the right side enriched region and the overall gradient seen in the measured data. PLS and PLS Spatial shift the intensity upward and over smooth the spatial pattern, leading to overly uniform maps with weakened localization. DGAT under predicts the enriched region and distorts the spatial gradient, while GAT shows noisy textures with misplaced signals. GraphSAGE again under predicts and weakens spatial variation. Tree based models without spatial features tend to underestimate the enriched region, and their spatial variants recover the right side enrichment more clearly but still deviate from the measured contrast.

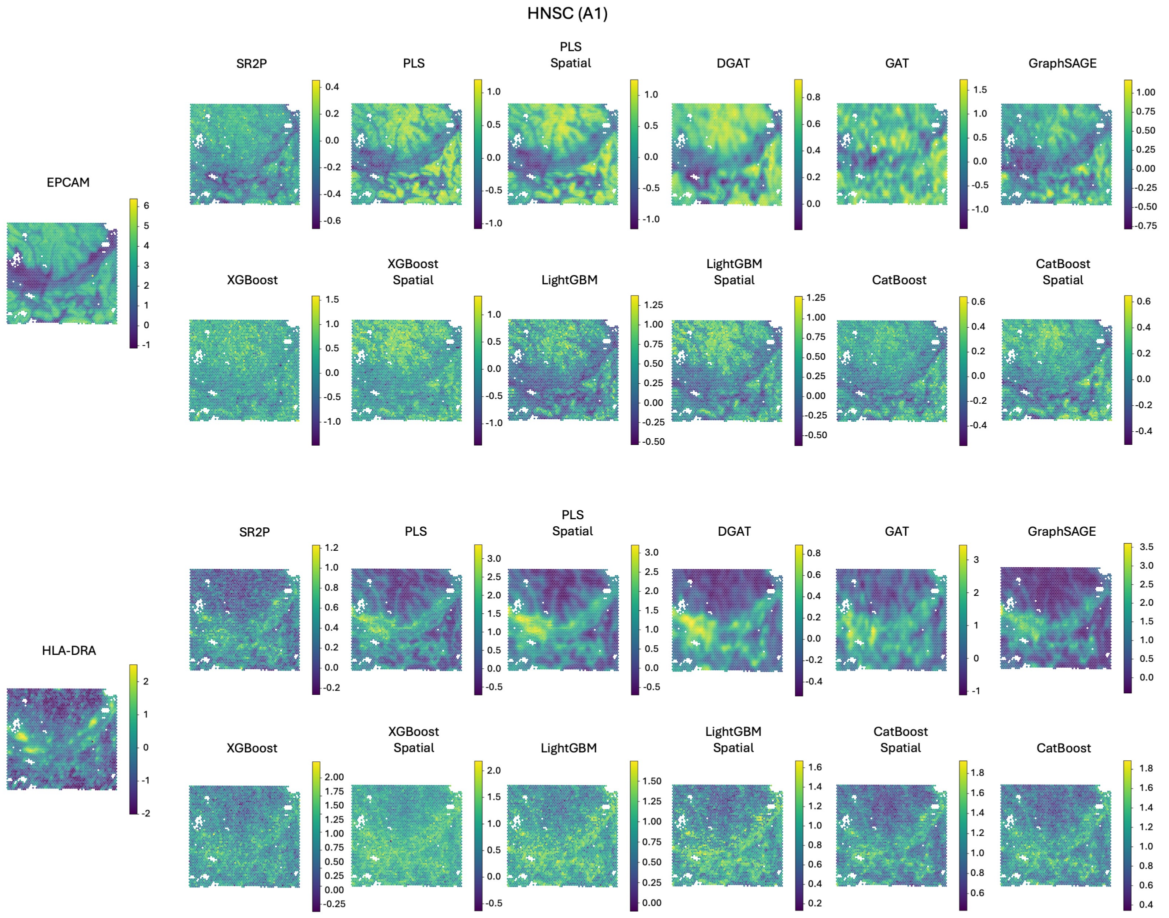


**Figure S12. Cross-tissue prediction of EPCAM and HLA-DRA expression in head and neck squamous cell carcinoma (HNSC, A1).** EPCAM is an epithelial cell marker and shows a structured spatial pattern in the measured data, characterized by regionally elevated expression with gradual transitions rather than sharp clusters. SR2P recovers the overall spatial layout and relative expression differences across regions, preserving the smooth transitions observed in the reference. In contrast, PLS and PLS Spatial substantially amplify EPCAM expression and introduce large high-expression areas that are not supported by the measured map. DGAT and GAT distort the spatial structure by producing patchy and fragmented patterns, while GraphSAGE shifts the overall intensity and weakens regional contrast. Tree-based models yield more stable predictions; however, XGBoost and LightGBM tend to introduce granular noise, and CatBoost, although more conservative, underestimates regional differences compared with the reference. HLA-DRA is an immune-related marker with localized enrichment superimposed on a heterogeneous background. SR2P captures both the location and relative intensity of these enriched regions, closely matching the measured spatial distribution. PLS based models exaggerate high-expression regions and reduce background heterogeneity, leading to inflated contrast. GNN based methods, particularly GAT and GraphSAGE, generate overly smoothed or mislocalized patterns that fail to align with the reference. Tree-based models partially recover enriched regions, with spatial variants improving continuity, but they frequently overestimate expression levels and introduce spurious local fluctuations.

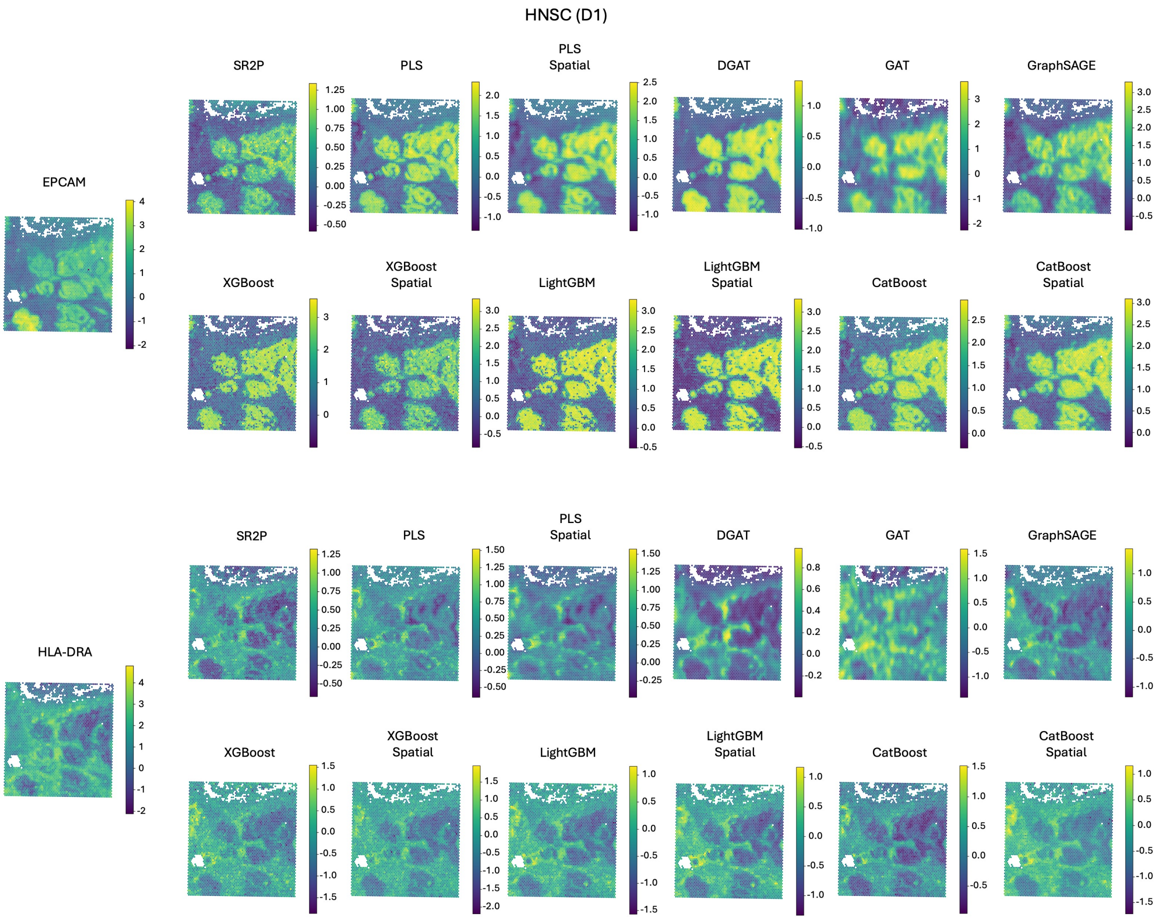


**Figure S13. Cross-tissue prediction of EPCAM and HLA-DRA expression in head and neck squamous cell carcinoma (HNSC, D1).** EPCAM is an epithelial marker and, in the D1 sample, exhibits clearly separated high-expression regions with sharp boundaries in the measured data, forming a block-like spatial pattern. SR2P closely matches both the location and shape of these epithelial regions, preserving boundary sharpness and relative intensity differences. PLS and PLS Spatial strongly amplify EPCAM expression and expand high-expression regions beyond their measured boundaries, leading to inflated epithelial areas. GNN-based methods show different failure modes: DGAT smooths regional boundaries and merges adjacent blocks, while GAT and GraphSAGE introduce intensity distortion and spatial artifacts that disrupt the measured block structure. Tree-based models recover the coarse epithelial layout; however, XGBoost and LightGBM introduce granular noise, and CatBoost tends to overestimate expression intensity, particularly in spatial variants. HLA-DRA is an immune-related marker with heterogeneous expression across the tissue. SR2P preserves the spatial heterogeneity and relative enrichment patterns observed in the measured data, without collapsing them into large uniform regions. In contrast, PLS-based models reduce heterogeneity and shift expression toward higher background levels. DGAT underestimates expression in enriched regions, while GAT and GraphSAGE produce patchy and spatially inconsistent patterns. Tree-based models partially recover heterogeneous signals, but both non-spatial and spatial variants tend to smooth local variation and distort relative intensities.

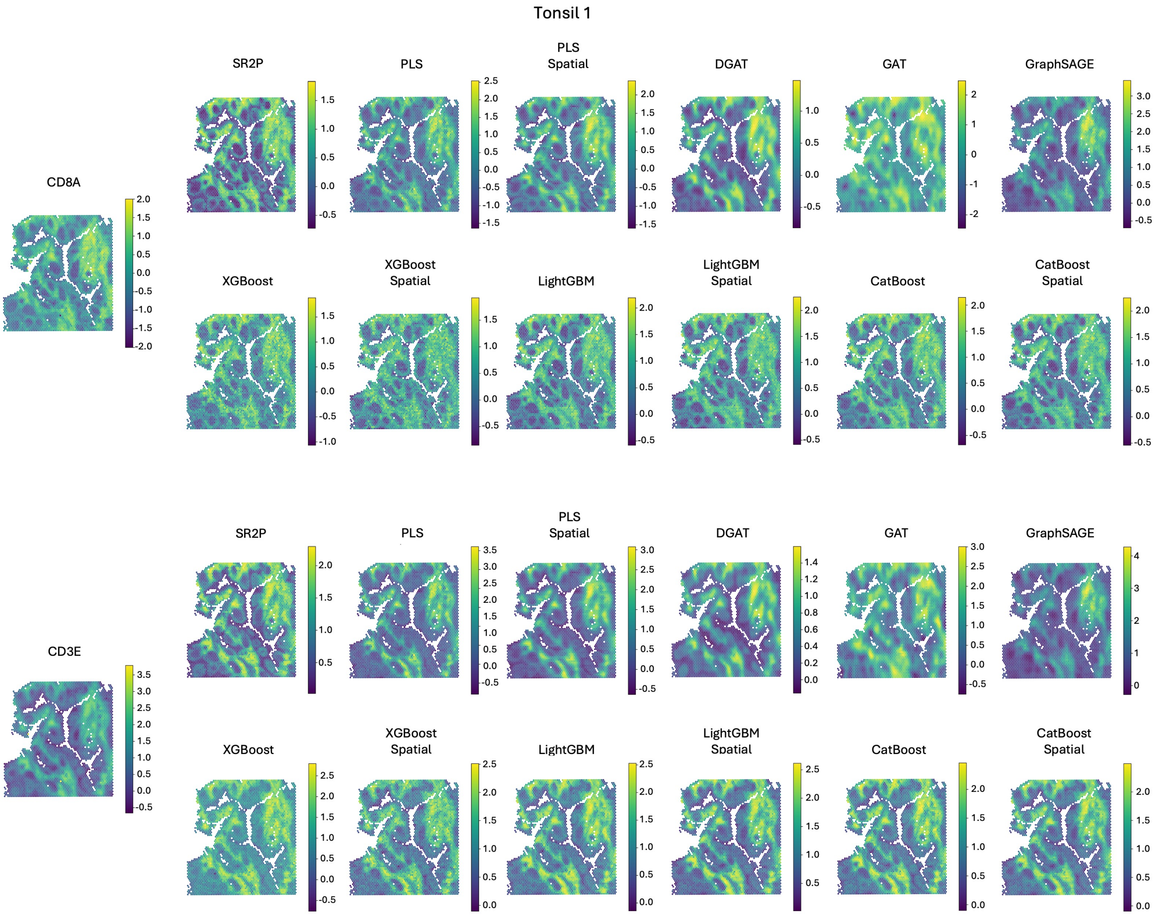


**Figure S14. Cross-tissue prediction of CD8A and CD3E expression in human tonsil (sample 1).** CD8A is a marker of cytotoxic T cells and displays a structured spatial pattern in the measured data, characterized by elongated high expression regions following lymphoid tissue architecture. SR2P preserves both the spatial continuity and the relative intensity of these CD8A enriched regions, maintaining clear separation between high and low expression areas. PLS and PLS Spatial exaggerate CD8A expression and expand high expression regions into surrounding tissue, reducing alignment with the measured boundaries. DGAT and GAT distort the elongated structures by introducing fragmented and patchy patterns, while GraphSAGE shifts the overall intensity and weakens contrast between immune enriched and depleted regions. Tree based models recover the general layout of CD8A expression, with spatial variants improving continuity; however, local intensity levels and boundary definition remain inconsistent with the reference. CD3E is a pan T cell marker with broader spatial distribution than CD8A, forming large immune enriched compartments in the measured data. SR2P accurately reproduces the extent and shape of these compartments, while preserving gradual intensity transitions. In contrast, PLS based models inflate expression levels and reduce spatial specificity. GNN based methods generate overly smoothed or spatially noisy predictions that blur compartment boundaries. Tree based models partially capture compartment structure, but both non spatial and spatial variants tend to over smooth local variation and alter relative expression intensity.

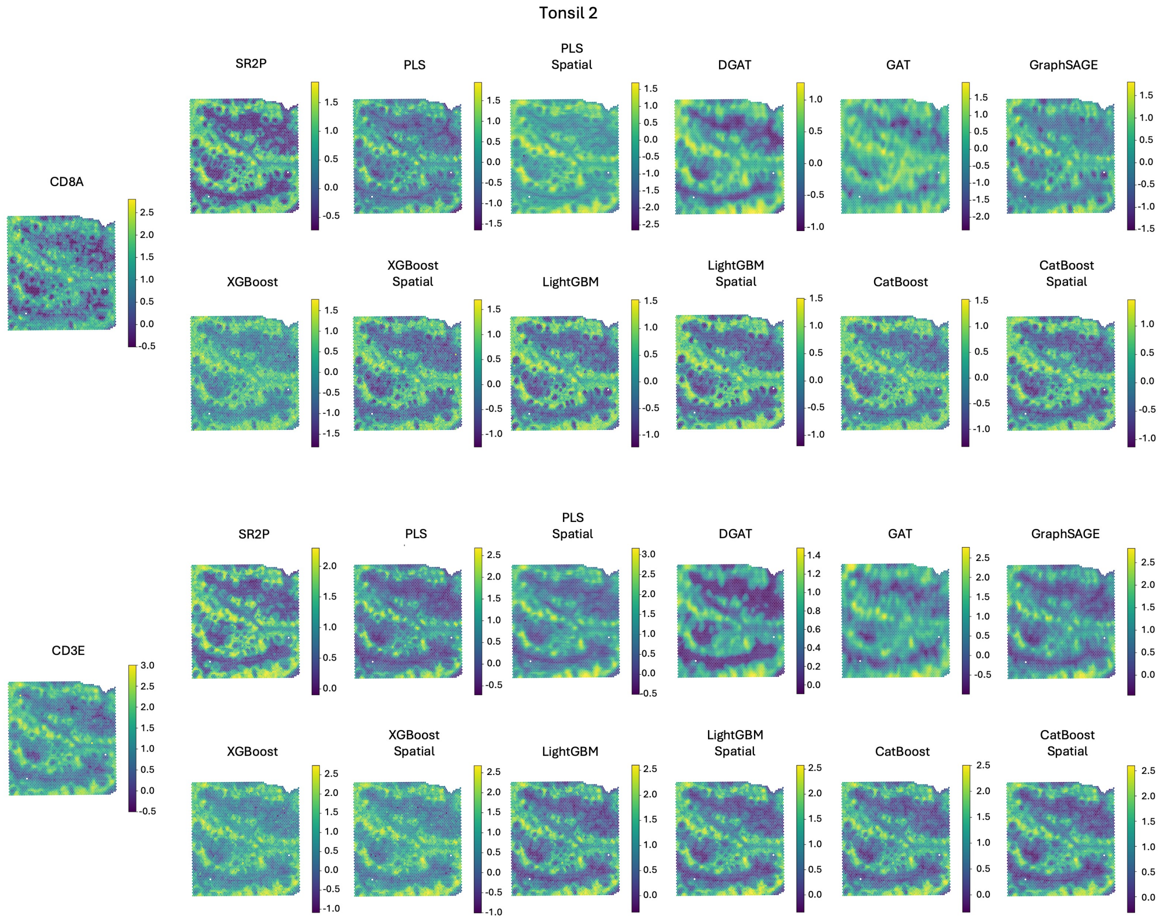


**Figure S15. Cross-tissue prediction of CD8A and CD3E expression in human tonsil (sample 2).** CD8A marks cytotoxic T cells and, in Tonsil 2, shows a more diffuse and finely patterned spatial distribution compared with Tonsil 1, with weak regional enrichment embedded in a heterogeneous background. SR2P preserves this diffuse spatial structure and maintains local variation without collapsing the signal into large uniform regions. In contrast, PLS and PLS Spatial alter the dynamic range and introduce broad intensity shifts, resulting in exaggerated high-expression regions that are not supported by the measured data. DGAT underestimates local variation and produces overly smoothed maps, while GAT and GraphSAGE introduce spatial noise and distort fine-scale patterns. Tree-based models generate visually similar reconstructions but tend to either attenuate local contrast or introduce granular artifacts; spatial variants improve continuity but do not fully recover the measured heterogeneity. CD3E is a pan–T cell marker with broader expression and similar diffuse spatial characteristics in Tonsil 2. SR2P maintains the weak gradients and fine-scale spatial variation observed in the measured map, avoiding artificial compartmentalization. PLS-based models inflate expression levels and reduce spatial specificity, whereas DGAT produces localized intensity depressions that are absent in the reference. GNN-based methods show patchy patterns with inconsistent signal localization. Tree-based models partially capture the overall distribution but smooth out subtle spatial variation and alter relative expression levels.

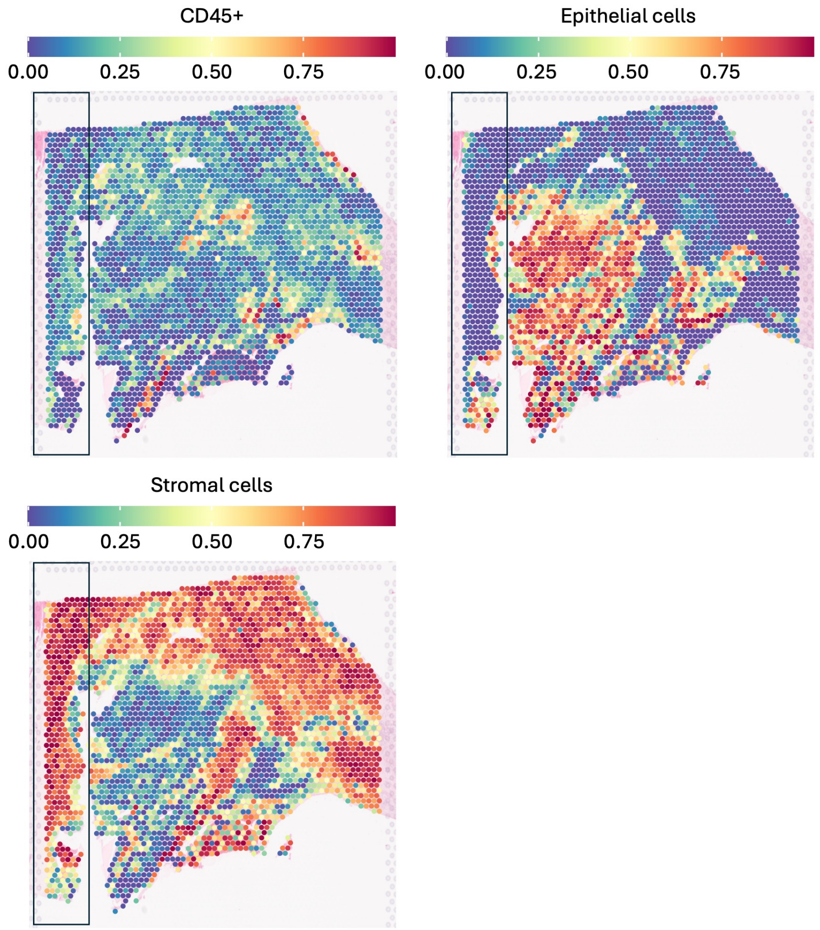


**Figure S16. Deconvolution maps for a poorly immune-infiltrated HNSCC sample.** Spatial maps of cell type proportions estimated by SCDC deconvolution are shown for representative immune and tissue compartments, including CD45+ immune cells, epithelial cells, and stromal cells. Each spot is color-mapped to the estimated proportion (0-1) and overlaid on the corresponding tissue image. The boxed region highlights an area where immune infiltration is weak and spatial boundaries are less distinguishable using transcriptomic signals alone, motivating the integration of predicted protein features to recover previously undercharacterized macrophage-enriched domains.
